# *Arabidopsis* SWR1-associated protein methyl-CpG-binding domain 9 is required for histone H2A.Z deposition

**DOI:** 10.1101/657296

**Authors:** Magdalena E. Potok, Yafei Wang, Linhao Xu, Zhenhui Zhong, Wanlu Liu, Suhua Feng, Bilguudei Naranbaatar, Shima Rayatpisheh, Zonghua Wang, James A. Wohlschlegel, Israel Ausin, Steven E. Jacobsen

## Abstract

Deposition of the histone variant H2A.Z by the SWI2/SNF2-Related 1 chromatin remodeling complex (SWR1-C) is important for gene regulation in eukaryotes, but the composition of the *Arabidopsis* SWR1-C has not been thoroughly characterized. Here identify interacting partners of a conserved *Arabidopsis* SWR1 subunit, ACTIN-RELATED PROTEIN 6 (ARP6). We isolated nine predicted components, and identified additional interactors implicated in histone acetylation and chromatin biology. One of the novel interacting partners, methyl-CpG-binding domain 9 (MBD9), also strongly interacted with the Imitation SWItch (ISWI) chromatin remodeling complex. MBD9 was required for deposition of H2A.Z at a distinct subset of ARP6-dependent loci. MBD9 was preferentially bound to nucleosome-depleted regions at the 5’ ends of genes containing high levels of activating histone marks. These data suggest that MBD9 is a SWR1-C interacting protein required for H2A.Z deposition at a subset of actively transcribing genes.

## Introduction

Chromatin dynamics plays a central role in regulating gene expression and has important implications in various cellular processes. The basic unit of chromatin is the nucleosome, which consists of 147 base-pairs (bp) of DNA wrapped around a histone octamer composed of two of each of the histones H2A, H2B, H3, and H4^1^. Cells contain chromatin remodeling complexes, which modulate the packaging, spacing, and composition of nucleosomes^2, 3^. The activity, targeting, and composition of remodeling complexes is tightly regulated to enable proper development of an organism^2, 3^.

The most highly conserved histone variant in eukaryotes is H2A.Z^4–6^. H2A.Z is found at many genes, most often around transcription start sites (TSS), with a particular preference for the +1 nucleosome^7–10^. The relationship between H2A.Z accumulation and transcription is complex. When present in the active genes of some organisms, the levels of H2A.Z at the TSS positively correlate with gene expression, typically when found together with H3/H4 acetylation, marks of active transcription^10–13^. H2A.Z is nonetheless absent at the most highly expressed genes^7^ and is also localized around the TSS of repressed genes that are poised for activation in yeast and animals^9, 14–16^. Although the function of gene body methylation is unclear, one study in *Arabidopsis* suggests that a function of gene body DNA methylation may be to prevent H2A.Z deposition over housekeeping genes^7, 17^, while another study found no such evidence^18^. However it is clear that an anti-correlation between H2A.Z and DNA methylation exists across diverse systems^16, 19–21^. In *Arabidopsis*, genes bearing high levels of gene body DNA methylation tend to be long and highly expressed, and show H2A.Z enrichment at their 5’-ends overlapping with activating histone modifications^17, 22–24^. By contrast, enrichment of H2A.Z over gene bodies (including high H2A.Z levels at the TSS) is typically found in shorter, weakly-expressed genes involved in environmental/stress responses that are marked with H3K27me3^17, 22–25^. Recently, levels of H3K27me3 were shown to be dependent on H2A.Z deposition in *Arabidopsis*^23, 26^. Together, these studies demonstrate that H2A.Z can participate in both activation and repression of gene expression, likely in concert with other factors and chromatin marks.

H2A.Z is deposited by the SWI2/SNF2-Related 1 (SWR1) complex, which involves the replacement of H2A-H2B dimers by H2A.Z-H2B dimers^27–29^. In yeast, the SWR1 complex (SWR1-C) is composed of 14 subunits^30^. In mammals, the structurally and functionally equivalent homolog of the yeast SWR1 complex, Snf2 Related CREBBP Activator Protein (SRCAP), is composed of 11 subunits, which are also conserved in yeast^31–33^. Based on homology, *Arabidopsis* contains one or several orthologs of the 11 conserved subunits from the yeast and human SWR1/SRCAP complexes, respectively^34^. Based on genetic and protein-protein interaction studies, several components have been validated as components of the *Arabidopsis* SWR1 complex, including PHOTOPERIOD-INDEPENDENT EARLY FLOWERING 1 (PIE1), the ATPase/scaffold subunit; ACTIN-RELATED PROTEIN 6 (ARP6); SERRATED LEAVES AND EARLY FLOWERING (SEF), the homolog of the yeast SWC6; and SWR1 complex subunit 2 (SWC2)^35–37^. A recent pull-down study of the SWC6 subunit identified additional subunits of the *Arabidopsis* SWR1 complex in yeast and validated their interaction with SWR1 complex subunit 4 (SWC4)^38^. Mutations in the genes encoding the SWR1 subunits share various phenotypes, with null mutations in the core catalytic subunit *pie1* producing the most severe phenotypes, including reduced size, loss of apical dominance, reduced fertility, and early flowering^35–37, 39–41^.

How the SWR1 complex is targeted across the genome remains unclear. Evidence suggests that H2A.Z localization around TSS in yeast is dependent on open chromatin and histone acetylation deposited by the Nucleosome acetyltransferase of H4 (NuA4) complex^8, 42–46^. The SWR1-C and NuA4 complexes share four common members: SWC4, YEAST ALL1-FUSED GENE FROM CHROMOSOME 9 (YAF9), ACTIN-RELATED PROTEIN 4 (ARP4), and ACTIN 1 (ACT1)^27–29, 47–49^. Homologs of these four proteins are present in the mammalian TIP60 complex, which possesses the dual activities of acetylating histones and depositing H2A.Z^31, 50, 51^. The *Arabidopsis* genome also contains homologs for many of the yeast NuA4 subunits; however, a functional link between histone acetylation and H2A.Z deposition is unclear^52^. YAF9 proteins are required for both the proper expression of *FLOWERING LOCUS C* (*FLC)* and H2A.Z acetylation at its regulatory regions, but not for H2A.Z deposition^53^. Much more needs to be learned about the potential NuA4 complex in *Arabidopsis*, including its composition and genome-wide effects on H2A.Z deposition and modification. Although genome-wide localization of the SWR1 components is lacking in *Arabidopsis*, it has been reported that ARP6 and SWC4 directly bind their target locus *FLOWERING LOCUS T* (*FT)*^38^; likewise, ARP6 and SWC6 bind directly to *FLC*^35^.

In this study, we used the conserved ARP6 subunit as bait to analyze the composition of the *Arabidopsis* SWR1 complex. We identified *Arabidopsis* homologs of most of the conserved SWR1 complex components in yeast, as well as several additional interacting partners: MBD9, four of the seven members of the ALFIN1-LIKE family (AL4, AL5, AL6, AL7), and two components of the yeast NuA4 and *Arabidopsis* SAGA histone acetyltransferase complexes, TRANSCRIPTION–ASSOCIATED PROTEIN 1A AND 1B (TRA1A and TRA1B). Our data are consistent with a related study where ARP6-interacting proteins were identified using ARP6 TAP-tag purification followed by mass spectrometry (Sijacic P et al, 2018, bioRxiv). We focused on characterizing MBD9, a protein implicated in flowering time regulation^54^. Using biochemical, genetic, and transcriptome analyses, we showed that MBD9 interacts with the SWR1 complex in *Arabidopsis*, and is required for H2A.Z deposition at a subset of *ARP6-*dependent genes. Furthermore, immunoprecipitation of MBD9 followed by mass spectrometry (IP–mass spec) revealed its association with the ISWI chromatin remodeling complex, suggesting additional functions for MBD9. Genome-wide ChIP-seq analysis of MBD9 showed that it localizes to nucleosome-depleted regions of active genes but does not influence open chromatin itself. MBD9 is thus a novel interacting partner of the *Arabidopsis* SWR1 complex required for H2A.Z deposition at a subset of actively transcribed genes.

## Results

### MBD9 interacts with the SWR1 complex and controls flowering timing

To identify components of the SWR1 complex in *Arabidopsis*, we used the conserved ARP6 subunit of the SWR1 complex as bait. We generated the epitope-tagged ARP6 lines *pARP6:ARP6-3xFLAG* and *pARP6:ARP6-9xMYC*. Both lines complemented *arp6-1* mutants (Supplementary Fig. 1), indicating that epitope-tagged ARP6 is functional *in vivo*. To detect as many potential protein interactions as possible, we performed IP– mass spec in both FLAG- and MYC-tagged ARP6 lines using highly proliferating inflorescence tissue where levels of proteins are high. In all replicates (three biological replicates with two technical replicates each), we observed enrichment of the *Arabidopsis* homologs of most of the conserved yeast SWR1 complex components---ARP6, PIE1, RuvB-like protein 1 and 2 (RIN1, RIN2), ARP4, SWC4, SWC2, YAF9A, and SEF. This includes three of the four common subunits of the yeast NuA4 complex, ARP4, SWC4, and YAF9A (Table 1). Our experiment confirms previous well-characterized *in vivo* and *in vitro* interactions of ARP6 with PIE1 and SEF^36–38^. We additionally identified two homologs of the yeast Tra1 proteins, subunits of the yeast NuA4 complex, TRA1A and TRA1B. We also observed consistent enrichment for several interacting partners previously implicated in chromatin biology, MBD9^54, 55^ and AL4, AL5, AL6, and AL7, which recognize di- and trimethyl H3K4 histone marks^56^. The interaction between ARP6 and MBD9 was further validated by co-immunoprecipitation (co-IP) in *Arabidopsis* flowers using an antibody raised against MBD9 (Fig. 1a). We then generated epitope-tagged MBD9 lines, *pMBD9:MBD9-3xFLAG* and *pMBD9:MBD9-9xMYC*. Both lines complemented *mbd9-3* mutants (Supplementary Fig. 2), indicating that the epitope-tagged MBD9 is functional *in vivo*. We performed IP–mass spec using floral tissue from both FLAG- and MYC-tagged MBD9 lines. In both experiments we observed enrichment of the SWR1 complex components PIE1, RIN1, ARP4, RIN2, SWC2, ARP6, SWC4, YAF9A, and SEF (Table 2). Interestingly, we also identified TRA1A and TRA1B, components of the NuA4 complex, as common interacting partners between ARP6 and MBD9. Additionally, in our MBD9 IP–mass spec we detected high enrichment of ISWI chromatin remodeling complex members that were not present among the ARP6-associated proteins.

**Figure 1.**
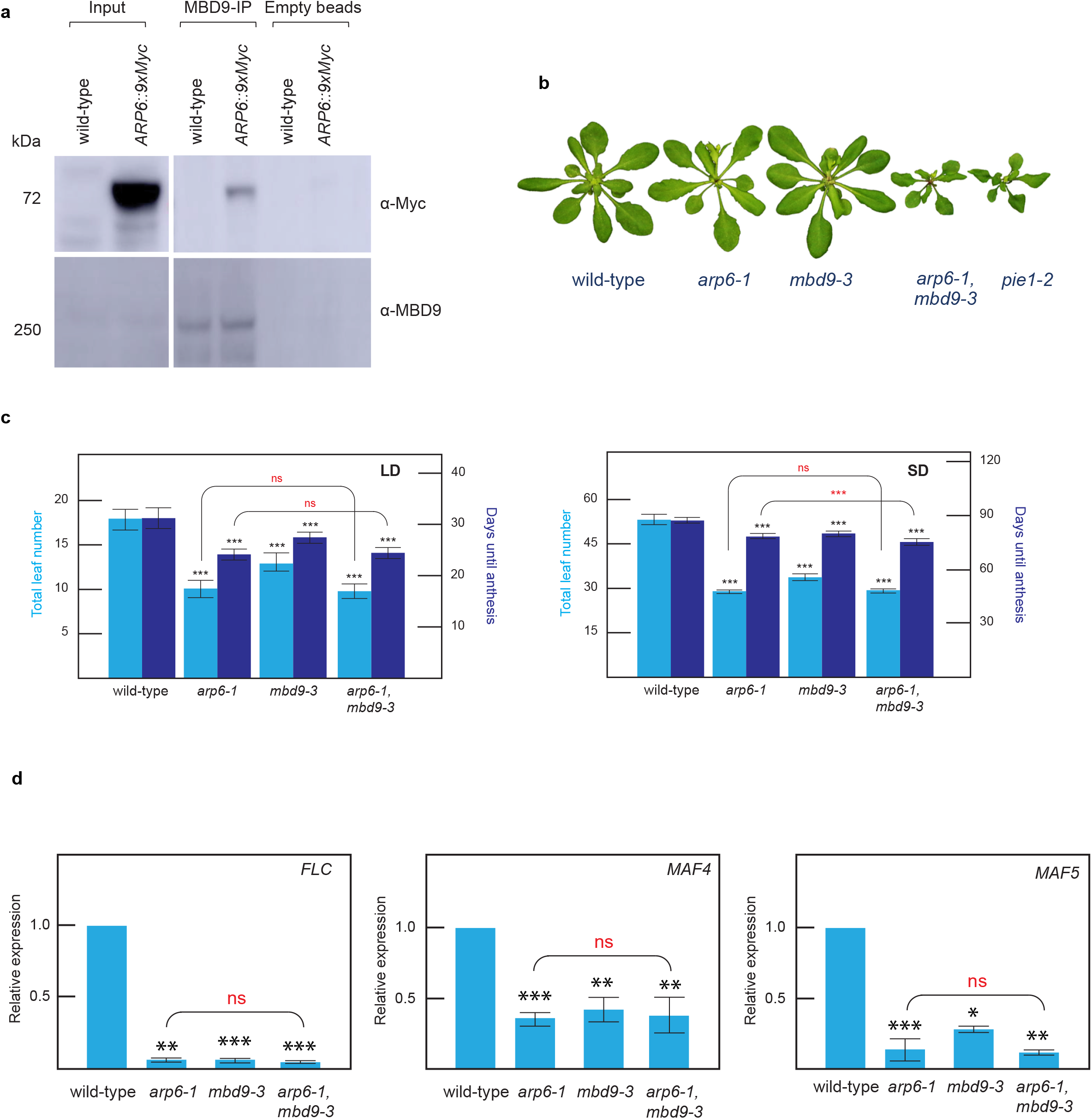
Interaction between ARP6 and MBD9 and phenotype of the *arp6-1 mbd9-3* double mutant. **(a)** Pull-down and co-immunoprecipitation assay. Empty bead lanes serve as negative controls. For each western blot, the antibody used is indicated to the right. **(b)** Phenotype of *arp6-1*, *mbd9-3*, *arp6-1 mbd9-3*, and *pie1-2* mutants. Plants were grown for 4 weeks (**b**) or 5 weeks (**c**) under long-day (LD) conditions (16 h light, 8 h dark) and short-day (SD) conditions (8 h light, 16 h dark). Flowering time expressed as the number of days from germination to anthesis as well as the total number of leaves produced by wild type, *arp6-1*, *mbd9-3*, and *arp6-1 mbd9-3* under the same conditions are shown. Averages from 16 plants ± standard deviations are shown. Paired two-tailed Student’s t-test was used to determine significance between wild-type and mutants (black asterisks) or between mutants (red asterisks); ns p-value > 0.05, * p-value <= 0.05, ** p-value <= 0.01, *** p-value <= 0.001. **(d)** Relative expression levels of *FLC*, *MAF4*, and *MAF5* in wild-type, *arp6-1*, *mbd9-3*, and *arp6-1 mbd9-3*. *UBQ10* was used as an internal control for expression analyses. Average expression levels from three independent replicates ± standard deviations are shown. Paired two-tailed Student’s t-test was used to determine significance between wild-type and mutants (black asterisks) or between mutants (red asterisks); ns p-value > 0.05, * p-value <= 0.05, ** p-value <= 0.01, *** p-value <= 0.001.

**Table 1.** IP–mass spec of ARP6. Spectral counts for ARP6-9xMyc and ARP6-3xFlag lines representing enriched genes over wild-type.

|  |  | ARP6-9xMYC |  |  |  | wild-type |  |  |  | ARP6-3xFLAG |  | wild-type |  |
| --- | --- | --- | --- | --- | --- | --- | --- | --- | --- | --- | --- | --- | --- |
| **Protein** | **Annotation** | **R1** | **R2** | **R3** | **R4** | **R1** | **R2** | **R3** | **R4** | **R1** | **R2** | **R1** | **R2** |
| AT3G33520 | ARP6 | 123 | 119 | 138 | 123 | 0 | 0 | 0 | 0 | 573 | 501 | 0 | 2 |
| AT3G12810 | PIE1 | 175 | 208 | 206 | 202 | 0 | 0 | 0 | 0 | 76 | 82 | 3 | 0 |
| AT5G22330 | RIN1 | 176 | 189 | 224 | 196 | 3 | 4 | 0 | 0 | 62 | 56 | 5 | 8 |
| AT5G67630 | RIN2 | 115 | 136 | 175 | 146 | 3 | 2 | 2 | 0 | 51 | 41 | 7 | 8 |
| AT1G18450 | ARP4 | 75 | 76 | 64 | 56 | 8 | 3 | 5 | 2 | 35 | 37 | 9 | 20 |
| AT2G47210 | SWC4 | 45 | 46 | 33 | 39 | 0 | 0 | 0 | 0 | 30 | 25 | 0 | 0 |
| AT2G36740 | SWC2 | 37 | 32 | 35 | 34 | 0 | 0 | 0 | 0 | 14 | 11 | 0 | 0 |
| AT5G45600 | YAF9A | 25 | 19 | 20 | 21 | 0 | 0 | 0 | 0 | 13 | 12 | 0 | 0 |
| AT5G37055 | SEF | 11 | 9 | 9 | 8 | 0 | 0 | 0 | 0 | 9 | 8 | 0 | 0 |
| AT3G01460 | MBD9 | 67 | 71 | 97 | 108 | 0 | 0 | 0 | 0 | 10 | 11 | 0 | 0 |
| AT2G17930 | TRA1A | 84 | 94 | 117 | 117 | 0 | 0 | 0 | 0 | 31 | 25 | 0 | 4 |
| AT4G36080 | TRA1B | 72 | 78 | 91 | 91 | 0 | 0 | 0 | 0 | 19 | 17 | 0 | 2 |
| AT5G26210 | AL4 | 2 | 3 | 2 | 2 | 0 | 0 | 0 | 0 | 0 | 0 | 0 | 0 |
| AT5G20510 | AL5 | 12 | 16 | 15 | 14 | 0 | 0 | 0 | 0 | 3 | 4 | 0 | 0 |
| AT2G02470 | AL6 | 5 | 6 | 4 | 5 | 0 | 0 | 0 | 0 | 2 | 0 | 0 | 0 |
| AT1G14510 | AL7 | 6 | 3 | 5 | 6 | 0 | 0 | 0 | 0 | 0 | 0 | 0 | 0 |

**Table 2.** IP–mass spec of MBD9. Spectral counts for MBD9-9xMyc and MBD9-3xFlag lines representing enriched genes over wild-type.

| **Protein** | **Annotati**<br>**on** | **MBD9-9xMYC** | **MBD9-**<br>**3xFLAG** | **wild-type-**<br>**9xMYC** | **wild-type-**<br>**3xFLAG** |
| --- | --- | --- | --- | --- | --- |
| AT3G01460 | MBD9 | 355 | 607 | 0 | 0 |
| AT3G12810 | PIE1 | 75 | 49 | 0 | 0 |
| AT5G22330 | RIN1 | 33 | 26 | 0 | 10 |
| AT1G18450 | ARP4 | 31 | 24 | 0 | 9 |
| AT5G67630 | RIN2 | 29 | 22 | 0 | 13 |
| AT2G36740 | SWC2 | 17 | 0 | 0 | 0 |
| AT3G33520 | ARP6 | 16 | 5 | 0 | 0 |
| AT2G47210 | SWC4 | 12 | 9 | 0 | 0 |
| AT5G45600 | YAF9A | 5 | 6 | 0 | 0 |
| AT5G37055 | SEF | 2 | 0 | 0 | 0 |
| AT2G17930 | TRA1A | 128 | 76 | 0 | 0 |
| AT4G36080 | TRA1B | 120 | 74 | 0 | 0 |
| AT2G02470 | AL6 | 2 | 0 | 0 | 0 |
| AT5G18620 | CHR17 | 122 | 190 | 0 | 3 |
| AT3G06400 | CHR11 | 118 | 177 | 0 | 4 |

To explore the genetic interaction between MBD9 and ARP6, we used the T-DNA mutants *arp6-1* and *mbd9-3*^35, 39, 40, 54, 55^. Both single mutants displayed an early flowering time phenotype (Fig. 1c) as described previously^39, 40, 54^. The *mbd9-3* phenotype was less severe than that of *arp6-1*; *mbd9-3* plants were larger, with a leaf morphology that more closely resembled wild-type plants (Fig. 1b). The *mbd9-3* mutant plants also flowered slightly later than the *arp6-1* mutants under both long-day (LD) and short-day (SD) conditions (Fig. 1c). The *arp6-1 mbd9-3* double mutants had a more severe developmental phenotype than either of the single mutants, which resembled the strong phenotype of *pie1-2*, a mutant affecting the catalytic subunit of the SWR1 complex (Fig. 1b)^35, 37, 57^. The stronger phenotype of the *arp6-1 mbd9-3* double mutant was also observed in a related study (Sijacic P et al, 2018, bioRxiv). Despite the severe developmental phenotype, the double mutant had a similar flowering time to *arp6-1* under both LD and SD conditions; however, the time until anthesis (floral opening) was slightly shorter under SD growth conditions in the double mutant than the *arp6-1* mutant (Fig. 1c), most likely due to morphological defects present in the *arp6-1 mbd9-3* double mutant. These data suggest that, regarding flowering time, ARP6 is epistatic to MBD9 in both LD and SD conditions. Expression analysis of key flowering time genes, *FLC* and *MADS AFFECTING FLOWERING 4 AND 5* (*MAF4* and *MAF5*), indicated that both the single mutants and the double mutant had a similar reduction of expression of these genes (Fig. 1d). Together, these results indicate that MBD9 and ARP6 act together in the same genetic pathway controlling flowering time as part of the SWR1 complex.

### MBD9 is required for the deposition of H2A.Z

To investigate the role of MBD9 in H2A.Z deposition, we analyzed H2A.Z levels in *Arabidopsis* seedlings before the major developmental transition to flowering in order to minimize secondary effects in mutants. We performed western blot on total extracts, and observed the levels to decline in the mutants: wild-type >> *mbd9-3* > *arp6-1* > *arp6-1 mbd9-3* (Fig. 2a). To investigate the impact of H2A.Z loss on chromatin, we performed two replicates of H2A.Z ChIP-seq in wild-type, *mbd9-3, arp6-1*, and *arp6-1 mbd9-3*, with H3 ChIP-seq as a control. Pearson correlation indicated a considerable overlap between the ChIP-seq replicates (Supplementary Fig. 3a). As with H2A.Z protein levels, we detected a sequential decrease in H2A.Z signal over H2A.Z peaks in wild-type and over protein-coding genes as follows: wild-type > *mbd9-3* > *arp6-1* > *arp6-1 mbd9-3* (Fig. 2b). By contrast, H3 levels were unchanged over these regions (Supplementary Fig. 3b). Using the MACS2 peak caller for each replicate (q-value less than 0.01), we defined common H2A.Z differential regions for each genotype. We surveyed the gene annotations in each differential region and identified genes that lost H2A.Z in the mutants compared to wild-type: 2,594 for *mbd9-3*, 12,222 for *arp6-1*, and 15,226 for *arp6-1 mbd9-3*. GO term enrichment for the three groups of genes yielded similar results, including post-embryonic development and protein modification, indicating that broad classes of genes were impacted by the loss of H2A.Z in MBD9 and ARP6 mutants (Supplementary Fig. 3c).

**Figure 2.**
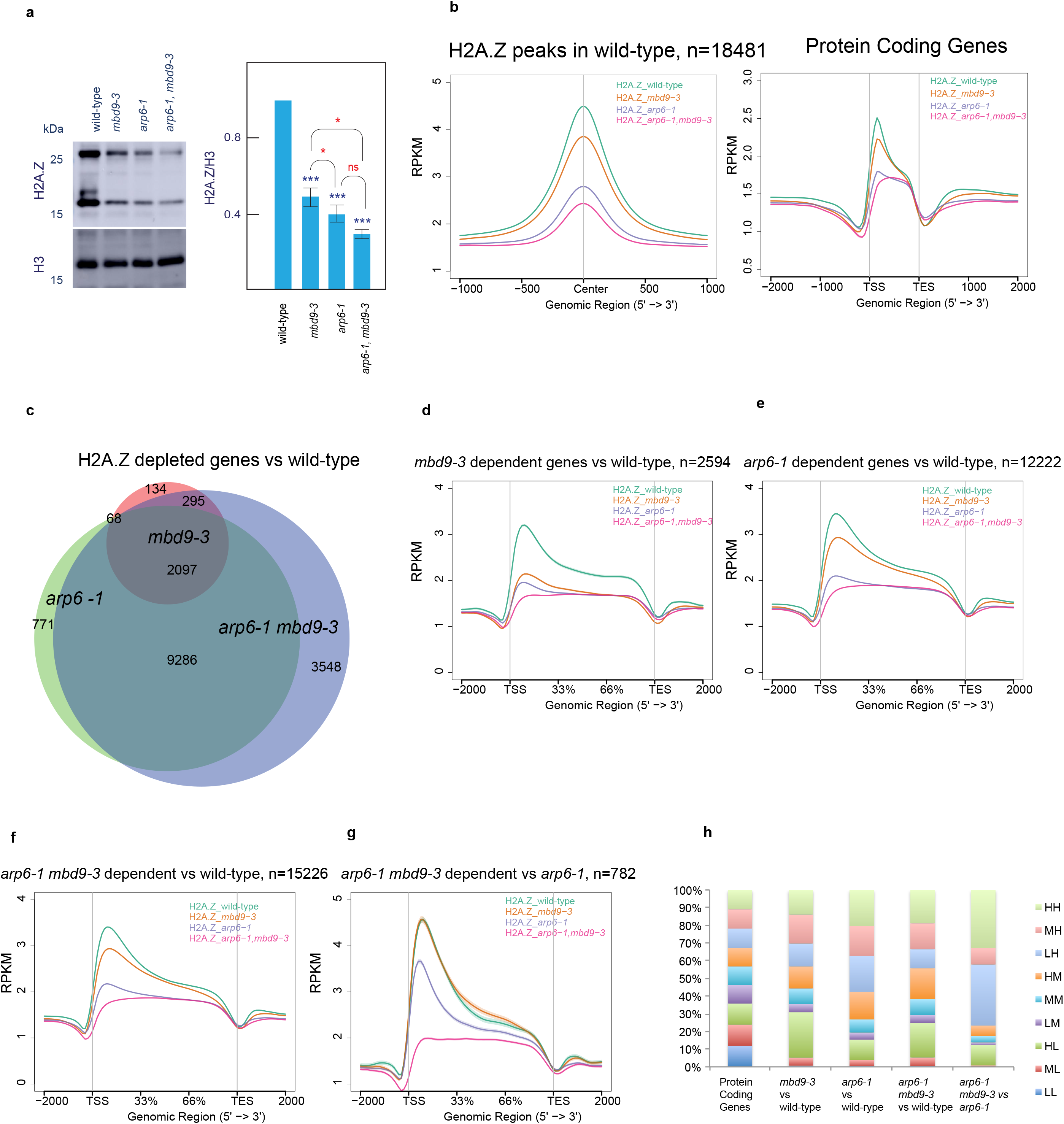
Characterization of H2A.Z in wild-type and mutant chromatin. **(a)** Western blot and quantification for H2A.Z and H3 levels in total extracts from wild-type, *mbd9-3, arp6-1*, and *arp6-1 mbd9-3*. The upper band in the H2A.Z blot likely represents a modified form of H2A.Z. Only the lower band was used in the quantification. Average levels of three independent westerns ± standard deviations are shown. Paired two-tailed Student’s t-test was used to determine significance between wild-type and mutants (black asterisks) or between mutants (red asterisks); ns p-value > 0.05, * p-value <= 0.05, ** p-value <= 0.01, *** p-value <= 0.001. **(b)** Distribution of normalized H2A.Z ChIP-seq signal as read count per kilobase per million mapped reads (RPKM) from merged replicates over H2A.Z common peaks in wild-type and protein-coding genes. **(c)** Overlap of H2A.Z-depleted genes for *mbd9-3, arp6-1*, and *arp6-1 mbd9-3* mutants. **(d-g)** Distribution of normalized H2A.Z ChIP-seq signal (RPKM) from merged replicates over H2A.Z enriched genes for indicated mutants. **(h)** Intersection of H2A.Z-depleted genes from (2d-g) with nine classes of genes characterized by their distribution of H2A.Z in wild-type at TSS and gene body. Labels H – high, M – medium, L-low, refer to levels of H2A.Z at TSS (first letter) and gene body (second letter).

Comparison of the H2A.Z-depleted genes in each mutant revealed a high degree of overlap (Fig. 2c). A metaplot of H2A.Z occupancy over the different groups of genes revealed that: 1) although *mbd9-3* mutants had fewer genes that lost H2A.Z than *arp6-1*, the depletion of H2A.Z over *mbd9-3*-dependent loci was nearly as great as in *arp6-1* (Fig. 2d); 2), genes that lost H2A.Z in *arp6-1* and *arp6-1 mbd9-3* behaved similarly, with H2A.Z enrichment following the order: wild-type > *mbd9-3* > *arp6-1* > *arp6-1 mbd9-3* (Fig. 2e,f). We also defined 782 genes that required both MBD9 and ARP6 for H2A.Z deposition (*arp6-1 mbd9-3* mutant vs *arp6-1*, qvalue less than 0.01). These genes were enriched for cell cycle progression functions (Supplementary Fig. 3c). The H2A.Z profiles over this set of genes (Fig. 2g) support the observation that these regions were dependent on both MBD9 and ARP6 for H2A.Z deposition. In *Arabidopsis*, housekeeping genes typically contain H2A.Z exclusively over the +1 nucleosome and lowly expressed/responsive genes contain H2A.Z over the entire gene bodies, with highest levels of H2A.Z at the TSS^17, 23, 24^. To investigate which types of genes are most impacted by the loss of H2A.Z in our mutants, we divided genes (into nine classes) according to the levels of H2A.Z in wild-type over gene bodies (low, medium, and high) and TSS (low, medium, and high) as described in Coleman-Derr, D. and Zilberman D., 2012 (Supplementary Fig. 4). We intersected these classes of gene with genes that lost H2A.Z in our mutants and plotted their distribution (Fig. 2h). This revealed that loss of H2A.Z in the *mbd9-3* mutant tends to occur over genes possessing high levels of H2A.Z over the TSS and low levels of H2A.Z over gene bodies (HL class– High TSS, Low gene body), whereas loss of H2A.Z in the *arp6-1* mutant tends to occur over genes containing high levels of H2A.Z over gene bodies (LH, MH, and HH classes) (Fig. 2h).

### MBD9/ARP6 partial redundancy in gene expression control

To investigate the relationship between H2A.Z loss and transcription, we performed RNA-Seq in seedlings in wild-type, *mbd9-3, arp6-1*, and *arp6-1 mbd9-3*. We defined differentially expressed genes (fold change greater than 2, FDR less than 0.05). We detected 277 mis-regulated genes in *mbd9-3*, 2,042 in *arp6-1*, and 1,620 in *arp6-1 mbd9-3*. Among the down-regulated genes we detected *FLC*, *MAF4*, and *MAF5* as targets of both MBD9 and ARP6, confirming our RT-qPCR results (Supplementary Fig. 5a). Importantly, the expression of genes encoding the predicted subunits of the SWR1 complex as well as H2A.Z genes were largely unaffected in the mutants (Supplementary Fig. 5b). We examined down-regulated and up-regulated genes (Fig. 3) and observed a high degree of overlap between the *arp6-1* and *arp6-1 mbd9-3* mutants, with 584 common up-regulated and 616 common down-regulated genes (Fig. 3a,b). These overlapping genes showed the same trend in expression change in *mbd9-3* mutants, but to a lesser degree that did not meet our significance cutoffs (Fig. 3c,d). We also defined genes specifically up- and down-regulated in either *mbd9-3* or *arp6-1*, implying independent functions at a subset of loci (Fig. 3a-d), as well as a set of 218 up-regulated genes that were dependent on both ARP6 and MBD9 for expression (Fig. 3a-d), suggesting a redundant role of MBD9 and ARP6 at this set of genes. By contrast, the unique down-regulated genes (n=170), did not show a significant change between *arp6-1* and *arp6-1 mbd9-3* (Fig. 3d). GO term analysis indicated that up-regulated genes in *mbd9-3* mutants are involved in post-embryonic development and response to stress, whereas for *arp6-1* and *arp6-1 mbd9-3* up-regulated genes are involved in lipid localization, biosynthetic processes, and response to drug (Supplementary Fig. 5c). By contrast, down-regulated genes are involved in response to stress, stimulus, and defense in all mutants (Supplementary Fig. 5c).

**Figure 3.**
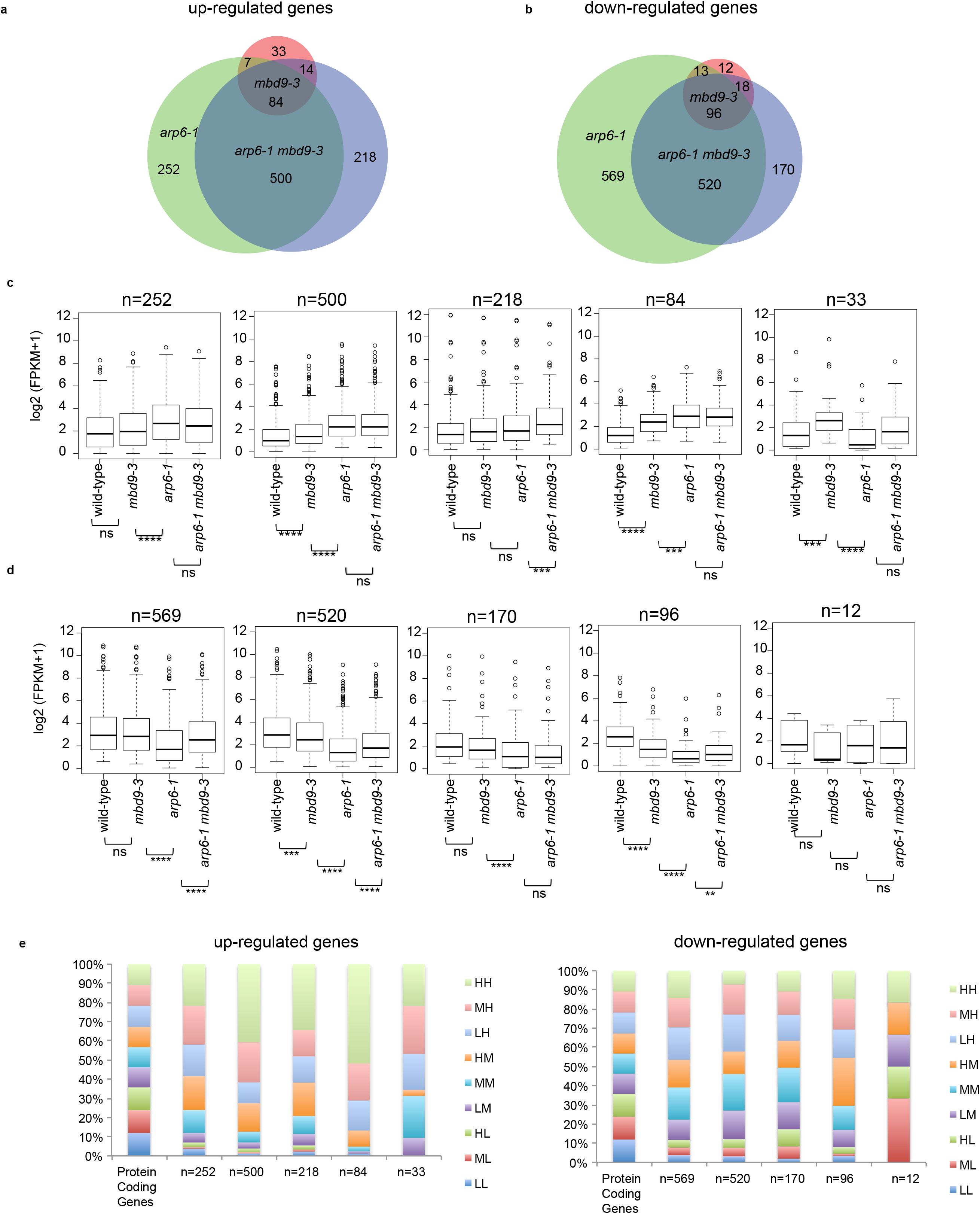
RNA-Seq and H2A.Z levels in **wild-type**, *mbd9-3, arp6-1*, and *arp6-1 mbd9-3* mutants. **(a)** Overlap between significantly (log_2_ >= 1, FDR <=0.05) up-regulated and **(b)** down-regulated genes for each mutant vs wild-type control. Boxplots of average normalized RNA-Seq reads (FPKM +1) represent different classes of overlapping and non-overlapping **(c)** up-regulated and **(d)** down-regulated genes based on intersection. Unpaired two-samples Wilcoxon test was used to determine significance: ns p-value > 0.05, * p-value <= 0.05, ** p-value <= 0.01, *** p-value <= 0.001, **** p-value <= 0.0001. (e) Intersection of up-regulated and down-regulated genes with nine classes of genes characterized by their distribution of H2A.Z in wild-type at TSS and gene body. Labels H – high, M – medium, L-low, refer to levels of H2A.Z at TSS (first letter) and gene body (second letter).

We compared differentially expressed genes in relation to H2A.Z changes in the mutants. First, to obtain general trends, we generated boxplots of expression and profiles of H2A.Z in the nine classes of genes divided according to their H2A.Z occupancy (Supplementary Fig. 4) for wild-type, *mbd9-3*, *arp6-1*, and *arp6-1 mbd9-3* (Supplementary Fig. 6a,b). One general trend stood out, genes containing high-medium H2A.Z levels over gene bodies and the highest H2A.Z levels over TSS showed increased expression upon loss of H2A.Z in *arp6-1*, which was even more pronounced in the *arp6-1 mbd9-3* mutant (Supplementary Fig. 6a,b). Only a weak loss of H2A.Z was observed in the *mbd9-3* mutant over these classes of genes where expression was not significantly increased (Supplementary Fig. 6a,b). Interestingly, looking at a typical H2A.Z profile of housekeeping genes (low H2A.Z at gene body with high levels of H2A.Z at TSS), H2A.Z deposition depended on both ARP6 and MBD9; however, this loss did not cause major transcriptional changes even in *arp6-1 mbd9-3* where H2A.Z levels were nearly depleted (Supplementary Fig. 6a,b). This perhaps is not surprising since genes that typically change in expression (become upregulated) upon H2A.Z misregulation are weakly expressed/environmentally responsive genes that contain highest levels of H2A.Z over the gene body and the TSS^17, 23–25, 58, 59^, most likely reflecting changes in chromatin accessibility^23, 25^. We next profiled H2A.Z changes in the significantly up- and down-regulated genes in our RNA-Seq data sets (Fig. 3e). Both up- and down-regulated genes lost H2A.Z in *arp6-1* and *arp6-1 mbd9-3* mutants (Supplementary Fig. 5d). In agreement with the observed general trends, the up-regulated genes tend to belong to gene classes containing high levels of H2A.Z over the gene body (LH, MH, HH classes). More specifically, loss of H2A.Z in *mbd9-3* was observed most over genes up-regulated in *mbd9-3* (n=84 and n=33) and those specifically upregulated in *arp6-1 mbd9-3* (n=218) (Supplementary Fig. 5d), suggesting that MBD9-dependent H2A.Z deposition may be directly involved in repression of this subset of genes. *arp6-1* dependent up-regulated genes (n=252 and n=500) had minimal if any change in H2A.Z levels in *mbd9-3* (Supplementary Fig. 5d). The down-regulated genes did not tend to belong to any particular class of H2A.Z-containing genes (Fig. 3e). Except for few well-studied genes (MADS-box proteins), the down-regulated genes in the SWR1-C mutants (including MBD9) most likely reflect indirect effects related to H2A.Z loss perhaps due to misregulated transcription factors in our mutants (Supplementary Fig. 7). Altogether, these data suggest that 1) *mbd9-3* has a much weaker effect on transcription than *arp6-1*; 2) genes misregulated in the *mbd9-3* background are a subset of genes misregulated in the *arp6-1* background; 3) MBD9 and ARP6 also cooperate to control the expression of a subset of up-regulated genes; and 4) although loss of H2A.Z is observed genome-wide in *arp6-1* and is even more pronounced in *arp6-1 mbd9-3*, genes that change in expression upon the loss of H2A.Z are generally those that contain high levels of H2A.Z at TSS and high-medium levels of H2A.Z over the gene body.

### MBD9 occupies nucleosome-depleted regions near active genes

To determine the localization of MBD9 in chromatin, we performed MBD9 ChIP-seq on our complementing *pMBD9:MBD9-3xFLAG* T4 transgenic line, with wild-type as a control (Supplementary Fig. 2). Using MACS2 (q-value less than 0.01), we defined 10,348 MBD9-occupied peaks, corresponding to 5,772 genes having a peak within 1 kb upstream of the TSS. The distribution of all MBD9 peaks showed a preference for promoter and TSS regions (65%) compared to a control set of peaks randomly selected from the genome based on length and number (Fig. 4a). Although MBD9 contains a methyl-CpG-binding domain, it is predicted not to bind methylated DNA since it lacks key conserved residues required for mCpG binding^60^. Consistently, the level of methylation over MBD9 peaks in wild-type plants was lower in all cytosine contexts than over a control set of peaks (Fig. 4b).

**Figure 4.**
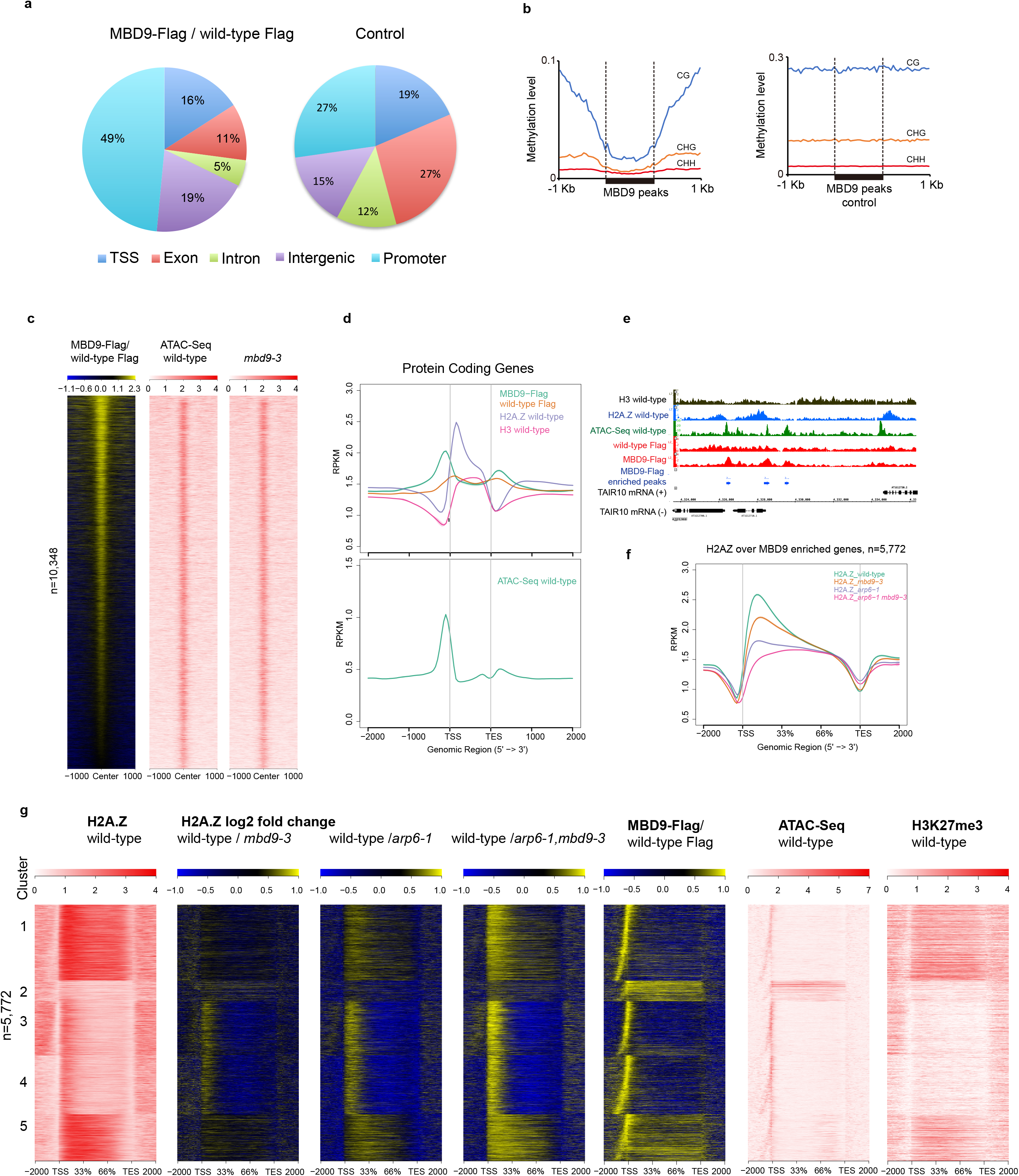
Characterization of MBD9 chromatin binding. **(a)** Genomic distribution of MBD9-enriched peaks vs wild-type (mac2 peak caller, q-value less than 0.01). **(b)** Average distribution of DNA methylation levels in CG, CHG, and CHH contexts in wild-type over MBD9-occupied peaks and MBD9 control peaks. **(c)** Heatmap of normalized MBD9 signal (log_2_ RPKM) over wild-type control and normalized ATAC-Seq signal (RPKM) from merged replicates in wild-type and *mbd9-3* mutant over MBD9-occupied peaks. **(d)** Upper: Distribution of normalized signal (RPKM) for MBD9-3xFlag ChIP-seq, wild-type Flag ChIP-seq control, H2A.Z, and H3 in wild-type over protein-coding genes. Lower: Distribution of normalized signal (RPKM) for ATAC-Seq in wild-type over protein-coding genes. **(e)** Snapshot representing normalized read counts (RPKM) for MBD9-3xFlag ChIP-seq, wild-type Flag ChIP-seq control, ATAC-Seq, H2A.Z, and H3. Signal for ATAC-Seq, H2A.Z, and H3 is a merge of two biological replicates. **(f)** Distribution of normalized H2A.Z ChIP-seq signal (RPKM) from merged replicates in wild-type, *mbd9-3, arp6-1*, and *arp6-1 mbd9-3* over MBD9-occupied genes. **(g)** Heatmap ordered based on *k*-means clustering of H2A.Z ChIP-seq signal in wild-type (cluster n=5) at MBD9-occupied genes. Shown are corresponding H2A.Z ratio plots for wild-type over *mbd9-3*, *arp6-1*, and *arp6-1 mbd9-3* mutants and MBD9-3X Flag ChIP-seq over wild-type flag control in log_2_ RPKM, ATAC-Seq in wild-type in RPKM, and H3K27me3 levels in wild-type in RPKM.

We plotted MBD9 peaks alongside open chromatin regions, as determined by ATAC-Seq, given MBD9’s clear preference for promoter regions. This revealed a significant overlap between MBD9 peaks and open chromatin regions (Fig. 4c). At these regions, the open chromatin was largely unaffected in *mbd9-3* (Fig. 4c). Metaplots of MBD9 ChIP-seq signal over protein-coding genes revealed two major MBD9 peaks, one over nucleosome-depleted regions just upstream of the TSS and a smaller peak just 3’ of the transcription end site (TES) (Fig. 4d). At the 5’ end, the MBD9 peaks overlap with the ATAC-Seq peaks and are just upstream of the H2A.Z peak (Fig. 4d,e).

We further investigated MBD9 occupancy in relation to H2A.Z distribution. When plotting H2A.Z over MBD9-occupied genes, we detected a loss of H2A.Z at the 5’-ends of genes in the *mbd9-3* mutant (Fig. 4f). To investigate whether the loss of H2A.Z occurs over a specific subset of genes, we ordered MBD9-occupied genes based on H2A.Z occupancy in the wild-type using *k*-means clustering (n=5) to reflect typical patterns of H2A.Z occupancy in *Arabidopsis*^17^ (Fig. 4g). This analysis revealed that MBD9 occupies genes with various H2A.Z localization patterns, including genes with low H2A.Z throughout (cluster 2, Fig. 4g) and genes containing H2A.Z over gene bodies (clusters 1 and 5, Fig. 4g). However, the locations that were most sensitive to the loss of H2A.Z in the *mbd9-3* mutants were those that contain H2A.Z predominately over their 5’-ends (clusters 3 and 4, Fig. 4g, ratio plots). These regions (clusters 3 and 4), contain low levels of H3K27me3. This is in contrast to the *arp6-1* and the *arp6-1 mbd9-3* mutants, where H2A.Z loss occurred throughout all MBD9-occupied genes, with the greatest loss observed in the double mutant (Fig. 4g).

### MBD9-enriched genes contain specialized chromatin features

To determine if specific chromatin features mark the regions that are enriched with MBD9, we analyzed ATAC-Seq data from wild-type plants and observed that MBD9-occupied genes tended to have more open chromatin than either an equivalent set of genes with similar expression levels, or all protein-coding genes (Fig. 5a,b). We also investigated preferences for histone modifications using previously published and newly generated data from equivalent tissues and stages of development. MBD9-occupied genes contained higher levels of H3K4me3 when compared to all protein-coding genes and the H3K4me3 signal was broader when compared to a control set of genes (Fig. 5c). A similar relative enrichment was also true for histone acetylation levels (Fig. 5c), whereas MBD9-occupied genes were depleted of H3K27me3 when compared to gene averages (Fig. 5c). This general trend was also true for regions depleted of H2A.Z in *mbd9-3*, as depicted in the heatmap ordered by expression level at all protein-coding genes (Supplementary Fig. 8). Regions that lost H2A.Z most in *mbd9-3* tended to occur predominantly at the 5’-ends of genes. These regions also contained high levels of activating histone modifications (H3K4me3, H3Ac) and low levels of H3K27me3, and coincided with MBD9-occupied peaks and open chromatin. In agreement, a related study also identified an enrichment of histone acetylation marks at MBD9-dependent H2A.Z sites (Sijacic P et al, 2018, bioRxiv).

**Figure 5.**
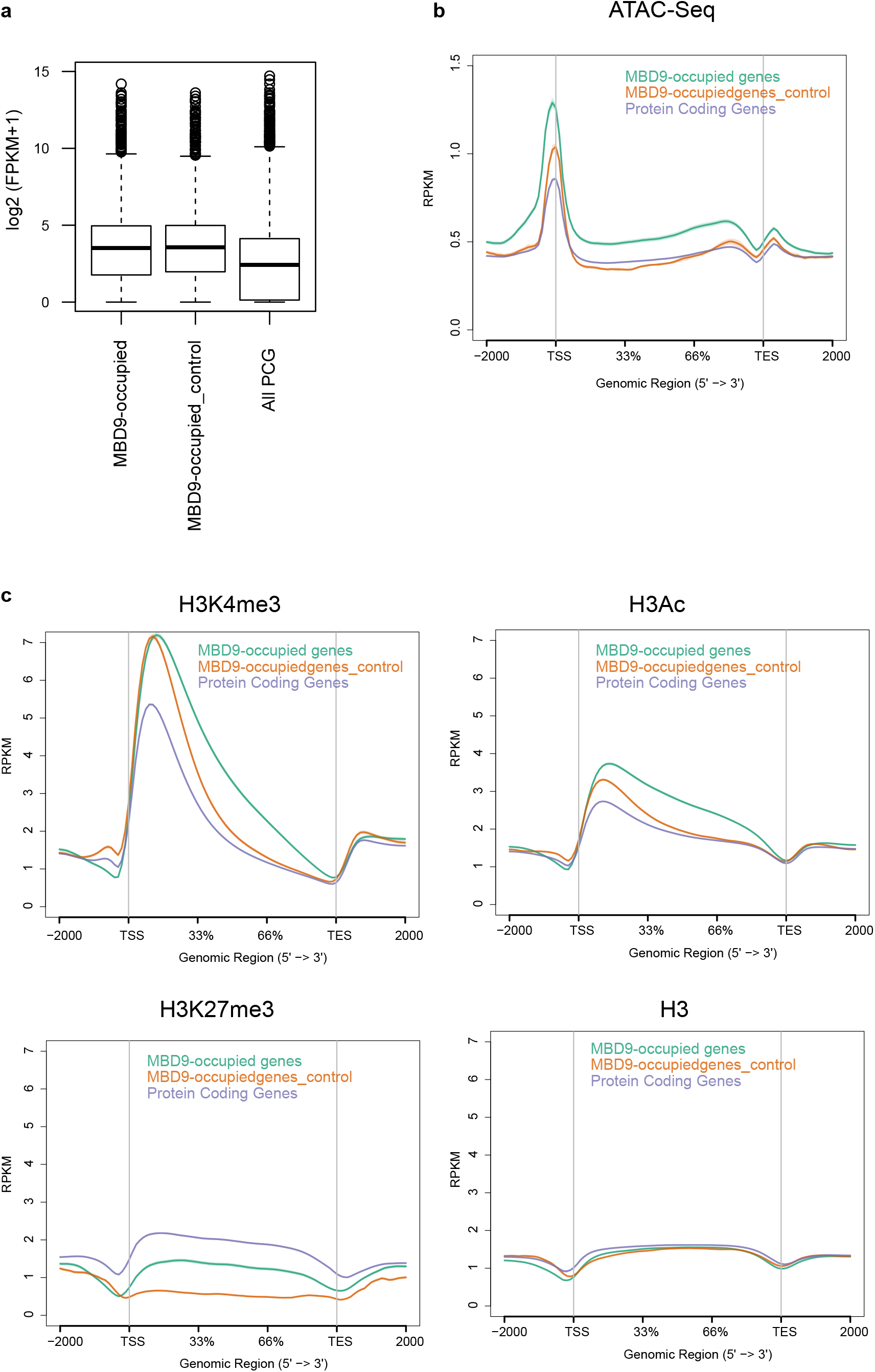
Chromatin features at MBD9-occupied genes. **(a)** Average expression levels log_2_ (FPKM +1) from RNA-Seq in wild-type for MBD9-occupied genes, MBD9 control genes, and protein-coding genes. **(b)** Distribution of normalized ATAC-Seq levels (RPKM) over MBD9-occupied genes, MBD9 control genes, and protein-coding genes. **(c)** Distribution of normalized levels (RPKM) for H3K3me3, H3Ac, H3K27me3, and H3 ChIP-seq in MBD9-occupied genes, MBD9 control genes, and protein-coding genes.

Altogether, these results suggest that MBD9 preferentially localizes to the open chromatin regions of actively transcribing genes, which contain high levels of H3K4me3 and histone acetylation and are different from genes controlled by the Polycomb Group Complex.

## Discussion

H2A.Z has been implicated in a wide variety of functions, including transcription, meiosis, and DNA damage. How it is specifically targeted to the locations where these events occur is poorly understood. In this study we have characterized the interacting partners of *Arabidopsis* ARP6, a homolog of Arp6 found in the yeast SWR1-C involved in H2A.Z deposition. We identified nine *Arabidopsis* proteins that are homologous to subunits common to the yeast (SWR1) and mammalian (SRCAP) complexes, ARP6, PIE1, RIN1, RIN2, ARP4, SWC4, SWC2, YAF9A, and SEF. We further identified several additional ARP6 interacting partners, TRA1A, TRA1B, MBD9, and four members of the Alfin1-like family (AL4, AL5, AL6 and AL7). These Alfin1-like containing proteins recognize H3K4me3 histones^56^, suggesting that they play a role in recruiting the SWR1 complex to H3K4me3 regions. The ARP6 interacting partners identified in our study are in agreement with a related study involving ARP6-TAP purification followed by mass spectrometry. However, there were a few differences. Specifically, we did not enrich for H2A.Z or ACT1 in our mass spec data, perhaps reflecting different methods of isolation, while we did identify TRA1A and AL4 as ARP6-interacting partners that were not reported by Sijacic P et al, 2018, bioRxiv.

We focused on characterizing the interaction between ARP6 and MBD9. We identified interacting partners for MBD9 and showed that it interacted with most components of the SWR1 complex, including PIE1, RIN1, ARP4, RIN2, SWC2, ARP6, SWC4, YAF9A, and SEF. MBD9 also interacted with TRA1A and TRA1B, further implicating histone acetylation in H2A.Z deposition. Additionally, we detected high enrichment of the ISWI chromatin remodeling complex in our MBD9 IP, which was not seen in the ARP6 IP–mass spec. The ISWI remodeling complex interacts with other chromatin remodelers^61^, MADS proteins^62^, and DTT-containing proteins^63^. MBD9 was amongst the reported interacting partners of CHR17, independently validating our results^61^. However, only two other SWR1 complex components (ARP4 and RIN1) were detected in the same CHR17 IP–mass spec study. Since we did not detect any interaction between CHR11/17 and ARP6 in our data, this suggests that MBD9 interacts with the two complexes separately. Our analysis of the genetic interactions between *mbd9-3* and *arp6-1* suggests that, despite the more severe developmental phenotype of the double mutant, ARP6 is epistatic to MBD9 regarding flowering time. These two observations are in agreement since flowering time is controlled by a particular set of genes, whereas H2A.Z deposition is globally affected in our mutants. The stronger developmental phenotype of the double mutant could have two plausible explanations: 1) multiple pathways are involved or 2) since the double mutant closely resembles *pie1-2* (a mutation affecting the catalytic subunit of the SWR1 complex), the phenotype results from MBD9 and ARP6 being part of the SWR1 complex, where the complex may be partly functional without one of the components but not without both. Future work will discriminate between these possibilities. Our genomic observations, H2A.Z ChIP-seq data, and transcriptome analysis revealed that MBD9 and ARP6 share many common targets, but the *mbd9-3* mutant shows a weaker phenotype than *arp6-1*. We observed that MBD9 is required for the deposition of H2A.Z at a subset of *arp6-1* dependent loci, similar to what was reported by Sijacic P et al, 2018, bioRxiv. We also identified a subset of genes that require both ARP6 and MBD9 for both their expression and for H2A.Z deposition, suggesting the two proteins act redundantly in some cases. Additionally, we determined the genome-wide occupancy of MBD9 and showed that it preferentially localized to the open chromatin regions of promoters and the transcription termination sites of active genes that have high levels of activating histone modifications (H3K4me3 and histone acetylation) and did not localize to genes repressed by the Polycomb Group Complex. These data suggest that the MBD9-containing SWR1 complex promotes H2A.Z deposition at the 5’ ends of highly expressed genes.

Regarding the interaction of MBD9 with the SWR1 complex, three possibilities exist based on our genetic and genomic observations: 1) MBD9 is always a member of the SWR1 complex but not always required, 2) MBD9 is part of a subset of all SWR1 complexes in *Arabidopsis*, 3) MBD9 only transiently interacts with the SWR1 complex. Based on size exclusion chromatography of ARP6-containing SWR1 complex in the *mbd9* mutant, Sijacic P et al, 2018, bioRxiv concluded that MBD9 is not a core subunit of the *Arabidopsis* SWR1 complex. However, as noted by the authors, this does not exclude the possibility that MBD9 is part of a small subset of all SWR1 complexes in *Arabidopsis*. Given that we detected most components of the SWR1 complex in our MBD9 IP–mass spec, our genetic and genomic data best fit the model that MBD9 is part of a subset of the SWR1-containing complexes.

Surprisingly, although we detected many genes that lost H2A.Z in the *mbd9-3* mutant, this loss did not result in major transcriptional changes. This may be because the loss of H2A.Z in the *mbd9-3* mutant occurs over genes with low H2A.Z over gene bodies and several studies suggest that genes changing most upon loss of H2A.Z are the ones bearing H2A.Z over the gene body^17, 23, 24^. One plausible explanation for the lack of transcriptional response at these regions in the *mbd9-3* mutant is redundancy. It is possible that redundant mechanisms are present at housekeeping genes to maintain their high expression and losing H2A.Z is not enough to perturb their activity. In support, H2A.Z is proposed to have redundant roles with chromatin remodeling complexes in yeast^64^. Interestingly, we identify components of the ISWI chromatin remodeling family as interacting partners of MBD9 in our IP. Additionally, CHR11/17 mutants have defects in flowering time and decreased levels of *FLC* expression^65^. Perhaps in addition to promoting H2A.Z deposition, MBD9 promotes low nucleosome occupancy at genes containing H2A.Z at +1 by interacting with chromatin remodelers to maintain these loci in a transcriptionally permissive state. In support, a recent study of FORGETTER1, a protein involved in heat shock memory response, was found to interact with chromatin remodelers including CHR11/17 to maintain low nucleosome occupancy of genes in non-inductive conditions^61^. In *Arabidopsis*, the ISWI remodeling complex is required for the regular spacing of nucleosomes downstream of the +1 border nucleosome^66^. Interestingly, many highly expressed genes in the CHR11/17 mutant have defects in nucleosome positioning, yet their transcription is unaffected^66^. This finding supports our hypothesis that redundant mechanisms maintain high expression of housekeeping genes. Higher order mutants between the SWR1 complex and chromatin remodelers are required to test this hypothesis.

In yeast, the SWR1 complex preferentially binds to nucleosome-depleted regions^42, 43^. Consistent with the role of MBD9 in the SWR1 complex, we found that MBD9 also localizes to nucleosome-depleted regions, in close proximity to H2A.Z deposited at the 5’-ends of gene bodies. These regions are devoid of DNA methylation, consistent with the prediction that the methyl-binding domain of MBD9 does not recognize methylated sequences^60^.

The genes bearing MBD9 also contain high levels of histone acetylation, consistent with the observation that histone acetylation in yeast promotes H2A.Z deposition. We also observed enrichment of three of the four common NuA4 histone acetylation complex subunits, SWC4, YAF9, and ARP4, in our IP–mass spec. We also identified two homologs of the yeast Tra1 proteins, TRA1A and TRA1B, in both ARP6 and MBD9 IPs. Tra1 is a component of the yeast NuA4 complex^50^. In *Arabidopsis*, TRA1B interacts with SWC4, a component of the predicted NuA4 complex^67^, and both TRA1A and TRA1B are components of the *Arabidopsis* SAGA acetyltransferase complex^68^. This suggests that various acetyltransferase complexes may be promoting H2A.Z deposition via histone acetylation. Since we do not detect the NuA4 scaffold or the catalytic subunit in our IP, it is unlikely that the ARP6-containing SWR1 complex represents a fusion of the *Arabidopsis* SWR1 and NuA4 complexes, similar to the mammalian TIP60 complex; rather, the TRA1A and TRA1B subunits may function to bridge the acetyltransferases with the SWR1 complex.

In summary, our study identified components of the SWR1 complex in *Arabidopsis* and shed light on the role of MBD9 in promoting H2A.Z deposition at a subset of highly-expressed genes. Further work addressing the function of the H2A.Z histone variant should reveal the nature of its dependency on factors such as histone acetyltransferase and chromatin remodeling complexes for carrying out its varied functions across the genome.

## Methods

### Plant materials

All *Arabidopsis* plants used in this study were of the Col-0 ecotype and were grown at 22 °C under long-day conditions (16 h light, 8 h dark). For the short-day flowering time experiment, plants were grown at 22 °C (16 h dark, 8 h light). The following *Arabidopsis* mutant lines were used: *mbd9-3* (SALK_039302), *arp6-1* (Sail_599_G03), and *pie1-2* (SALK_003776C).

### Quantitative real time PCR

Total RNA was prepared from whole seedlings (14 DAG) using the RNA Prep Pure Plant Kit (Tiangen Biotech, Beijing China). 2 µg of RNA was used for the preparation of cDNA using PrimeScript Reagent Kit with gDNA eraser (Takara, Japan). Signal detection, quantification and normalization were done using Quant Studio 6 proprietary Software (Life Technologies, USA).

### Co-immunoprecipitation

An anti-MBD9 polyclonal antibody was raised in rabbits using the 1,941—2,176 aa MBD9 peptide as an antigen. The antibody was coupled to magnetic beads using the Dynabeads antibody coupling kit (Invitrogen, USA) following the manufacturer’s recommendations. Approximately 2 g of floral tissue was ground in liquid nitrogen, 10 ml IP buffer (50 mM Tris pH 7.6, 150 mM NaCl, 5 mM MgCl_2_, 10% glycerol, 0.1% NP-40, 0.5 mM DTT, 1 µg/µl pepstatin, 1 mM PMSF, 1 protease inhibitor cocktail tablet (Roche, 14696200)) was added, the mix was incubated on ice for 20 min, and spun at 4,000 *g* for 10 min at 4 °C. The supernatant was filtered through a double layer of Miracloth. Anti-MBD9 antibody coupled to magnetic beads was added and incubated for 2 h at 4 °C with gentle rotation. Beads were washed six times with IP buffer, 5 min per wash at 4 °C with rotation, and then boiled. Western blotting and detection were performed with anti-Myc 9E10 mouse monoclonal antibody (Santa Cruz Biotechnology, sc-40).

### RNA-Seq and analysis

Total RNA was extracted from 13 DAG shoots grown on 1% MS supplemented with 1% sucrose under long-day conditions. Four replicates, grown on separate plates, were collected for each genotype. RNA was extracted using the Direct-zol RNA Miniprep kit (Zymo). For RNA-Seq, 1 µg of total RNA was used to prepare libraries using the TruSeq Stranded mRNA-Seq kit (Illumina). Reads were aligned to TAIR10 using Tophat^69^ by allowing up to two mismatches and mapping only to one location. FRPKM values and differential gene expression were analyzed using Cufflinks^69^ with default settings. GO term enrichment was determined using AgriGO^70^.

### Epitope-tagged transgenic ARP6 and MBD9 lines

Full-length genomic DNA fragments containing ∼1 kb (MBD9) or ∼1.5 kb (ARP6) of promoter sequence, together with genomic gene sequences up to the annotated/major stop codon of MBD9 and ARP6 were amplified using PCR (Supplementary Information). The PCR product was cloned into the pENTR/D vector (Invitrogen) and delivered into a modified pEG containing 3XFLAG and 9XMYC tags using the LR reaction kit (Invitrogen). The pEG destination vectors were transformed into Agrobacterium strain AGL0, followed by transformation using the floral dip method^71^.

### Affinity purification and mass spectrometry

Approximately 10 g of flowers from transgenic lines expressing T3 ARP6-3xFLAG or T4 MBD9-3xFLAG and from Col-0 plants as negative controls, were ground to a fine powder using a RETCH homogenizer (3 min @ 30 Hz/min) and suspended in 30 ml of IP buffer. Tissue was further dounce homogenized until lump-free, then centrifuged for 20 min at 4,000 *g* and 4 °C. The lysate was filtered through two layers of Miracloth. Supernatant was incubated with 200 µl anti-FLAG M2 magnetic beads (M8823, Sigma), at 4 °C for 2 h. The bead-bound complex was washed once with 10 mL IP buffer, then four times (5 min rotating at 4 °C with 1.5 ml IP buffer), followed by a final wash with IP buffer without NP-40. The FLAG-IP was eluted twice with 300 µl 250 µg/ml 3X FLAG peptides (Sigma, F4799) in TBS (50 mM Tris-Cl, pH 7.5, 150 mM NaCl), mixing for 15 min at 4 °C. The eluted protein complexes were precipitated by trichloroacetic acid and subjected to mass spectrometric analyses as previously described^72^. In the case of ARP6-9xMyc and MBD9-9xMyc, monoclonal 9E10 coupled to magnetic beads was used (88842, Pierce). The bead-bound complexes were washed six times with 1 mL of IP buffer. For each wash, the beads were rotated at 4 °C for 5 min. For ARP6-9xMyc, proteins were released from the Streptavidin beads by 3C cleavage overnight at 5 °C. 3C-GST was removed by incubation with GST couple magnetic beads (Pierce, USA). For MBD9-9xMyc, proteins were eluted twice with 100 µl 8 M Urea in 50 mM Tris pH 8.5, mixing for 15 min at 37 °C. The supernatant was TCA precipitated.

### Genome-wide ChIP sequencing and library generation

For histone modification ChIPs, about 0.5 g of shoots were collected from 13 DAG seedlings grown on 1% MS plates supplemented with 1% sucrose under long-day conditions. ChIPs were performed as described previously with minor modifications^73^. Samples were crosslinked *in vitro*, sheared using Bioruptor Plus (Diagenode). Libraries were generated with NuGEN Ovation Ultra Low System V2 kit, according to the manufacturer’s instructions, and were sequenced on an Illumina HiSeq 4000 instrument. H2A.Z ChIP assays were performed using polyclonal antibodies specific to *Arabidopsis* H2A.Zs HTA11 and HTA9. The antibodies were produced in rabbits and further purified using the peptides described in Deal, R.B., *et al.* 2007. Abcam1791 (Abcam) antibody was used for H3 ChIP assays and active motif 39140 (Active Motif) was used for pan-acetyl H3 ChIP assays.

The ChIP protocol for MBD9 is a variation of a previously published protocol^74^. About 5 g of 13 DAG T4 MBD9-3XFLAG and Col-0 seedlings grown on 1% MS were fixed *in vivo* using vacuum infiltration as follows. Protein-protein crosslinking was performed first using 1.5 mM EGS in 1X PBS for 10 min followed by three washes with PBS. Next, tissue was fixed with 1% formaldehyde in 1X PBS for 20 min. Crosslinking was stopped with 0.125 M glycine in 1X PBS and the seedlings were vacuum infiltrated for 5 min. Tissue was patted dry and frozen at −80 °C. Tissue was homogenized to a fine powder using a RETCH homogenizer (2 min @ 30 Hz/min) and resuspended with Nuclear Isolation Buffer (50 mM Hepes, 1 M Sucrose, 5 mM KCl, 5 mM MgCl_2_, 0.6% Triton X-100, 0.4 mM PMSF, 5 mM Benzamidine, 1x protease inhibitor cocktail tablet (Roche, 14696200). Lysate was filtered through one layer of Miracloth and centrifuged for 20 min at 2,880 *g* and 4 °C. The pellet was resuspended with 1 ml Extraction buffer 2 (0.25 M sucrose, 10 mM Tris-HCl pH 8, 10 mM MgCl_2_, 1% Triton X-100, 5 mM BME, 0.1 mM PMSF, 5 mM Benzamidine, and 1x protease inhibitor cocktail tablet (Roche, 14696200)), followed by centrifugation for 10 min at 12,000 *g* and 4 °C. The pellet was then resuspended in 500 µl Extraction buffer 3 (1.7 M sucrose, 10 mM Tris-HCl pH 8, 2 mM MgCl_2_, 0.15% Triton X-100, 5 mM BME, 0.1 mM PMSF, 5 mM Benzamidine, 1x protease inhibitor cocktail tablet (Roche, 14696200), layered over 500 µl extraction buffer 3 and centrifuged for 1 h at 12,000 *g* and 4 °C. The pellet was lysed with 400 µl Nuclei Lysis Buffer on ice (50 mM Tris pH 8, 10 mM EDTA, 1% SDS, 0.1 mM PMSF, 5 mM Benzamidine, 1x protease inhibitor cocktail tablet (Roche, 14696200). To lysed nuclei, 1.7 ml of ChIP Dilution Buffer (1.1% Triton X-100, 1.2 mM EDTA, 16.7 mM Tris pH 8, 167 mM NaCl, 0.1 mM PMSF, 5 mM Benzamidine, 1x protease inhibitor cocktail tablet (Roche, 14696200) was added and DNA was sheared on a Bioruptor Plus (Diagenode) (30 sec ON/ 30 sec OFF, Max power, 17 min at 4 °C). Sheared chromatin was centrifuged twice at max speed for 10 min at 4 °C. The supernatant was further diluted with ChIP dilution buffer up to 4 ml. 100 µl of sample was saved as input and the rest was split into 2 ml DNA LoBind tubes (Eppendorf, D1579630) and 10 ml of anti-Flag antibody was added (F1804, Sigma-Aldrich). After incubation overnight with rotation at 4 °C, 50 µl Dynabeads (equal mix of Protein A and G, Invitrogen 10004D/10002D) was added to chromatin and incubated for an additional 2 h. The magnetic beads were washed with 1 ml of the following buffers for 5 min rotating at 4 °C: 2X with Low Salt Buffer (150 mM NaCl, 0.2% SDS, 0.5% Triton X-100, 2 mM EDTA, 20 mM Tris pH 8), 1X High Salt Buffer (200 mM NaCl, 0.2% SDS, 0.5% Triton X-100, 2 mM EDTA, 20 mM Tris pH 8), 1X LiCl Wash Buffer (250 mM LiCl, 1% Igepal, 1% Sodium Deoxycholate, 1 mM EDTA, 10 mM Tris pH 8), and 1X with TE buffer (10 mM Tris pH 8, 1 mM EDTA). The immunocomplex was eluted from the beads twice with 250 µl elution buffer (1% SDS, 10 mM EDTA, 0.1 M NaHCO_3_), incubating for 20 min with shaking at 65 °C. 400 µl of elution buffer was added to the input samples. A total of 20 µl 5 M NaCl was added to each tube, the crosslink was reversed by incubation at 65 °C overnight. Residual protein was degraded with 20 µg Prot K in 10 mM EDTA and 40 mM Tris pH 8 at 45 °C for 1 h followed by phenol/chloroform/isoamyl alcohol extraction and ethanol precipitation. The pellet was washed with 70% EtOH and resuspended in 50 µl molecular grade water. Libraries were prepared using the NuGEN Ovation Ultra Low System V2 kit, according to the manufacturer’s instructions, and were sequenced on an Illumina HiSeq 4000 instrument.

### ChIP-seq analysis

ChIP-seq fastq reads were aligned to the TAIR10 reference genome with Bowtie^75^ using default settings and allowing only uniquely mapping reads. Duplicated reads were removed using SAMtools^76^. IGB genome browser was used to visualize the data and to generate snapshots^77^. Normalized read coverage tracks were generated using the USeq package Sam2Useq application^78^. ChIP-seq peaks in wild-type and mutants (Fig. 5B) were called by callpeak function in MACS2 (v2.1.1.)^79^. ChIP-seq data metaplots and *k*-means clustering were generated by NGSPLOT (v4.02.48)^80^. Enriched motifs in peak sets were identified using HOMER findMotifsGenome^81^. Annotation of peak locations were carried out using the HOMER annotatepeaks function^81^. GO term enrichment was determined using AgriGO^70^.

### ATAC-Seq

Inflorescence tissues were collected for nuclei collection as described previously^82^. ATAC-seq was performed as described previously with slightly modified nuclei collection^83^. Briefly, 5 g of inflorescence tissue was collected and immediately transferred into ice-cold grinding buffer (300 mM sucrose, 20 mM Tris pH 8, 5 mM MgCl_2_, 5 mM KCl, 0.2% Triton X-100, 5 mM β-mercaptoethanol, 35% glycerol). The samples were ground with Omni International General Laboratory Homogenizer at 4 °C and then filtered through a two-layer Miracloth and a 40-µm nylon mesh Cell Strainer (Fisher). Samples were spin filtered for 10 min at 3,000 *g*, the supernatant discarded, and the pellet resuspended with 25 ml of grinding buffer using a Dounce homogenizer. The wash step was repeated once and the nuclei resuspended in 0.5 ml of freezing buffer (50 mM Tris pH 8, 5 mM MgCl_2_, 20% glycerol, 5 mM β-mercaptoethanol). ATAC-seq read adaptors were removed with trim_galore and then mapped to the *Arabidopsis* thaliana reference genome TAIR10 using Bowtie (-X 2000 −m 1). Chloroplast and mitochondrial DNA aligned reads were filtered out and duplicate reads were removed using SAMtools^76^.

### Data availability statement

All high-throughput sequencing data generated in this study are available privately for the reviewers at NCBI’s Gene Expression Omnibus (GEO) and are accessible via GEO Series accession number GSE123263. The data will be made public after publication. Detailed bioinformatics analysis, primers used, and a list of publicly available data used in the paper are provided in the Supplementary Information file. The source data underlying Figs 1a, 1c, 1d, 2a, S1, S2, Tables 1, 2 and gene listsx underlying Figs 2c, 3, 4, and S6 are provided as a Source Data file.

## End Notes

### Acknowledgements

We thank Mahnaz Akhavan for support with high-throughput sequencing at the University of California, Los Angeles (UCLA) Broad Stem Cell Research Center BioSequencing Core Facility. Work in the Jacobsen lab was supported by NIH grant R35 GM130272. M.E.P. was supported by the Damon Runyon Cancer Foundation DRG:2217-15. The work in the I.A. laboratory was supported by the National Science Foundation of China (grant numbers: 31571328 and 31870270). S.E.J. is an Investigator of the Howard Hughes Medical Institute.

### Author Contributions

M.E.P. performed RNA-Seq and ChIP-seq and generated sequencing libraries. Y.W. and L.X. developed ARP6-tagged lines, performed IP, co-IP, western blots, RT-PCRs, and genetic analyses. M.E.P. and B.N. developed MBD9-tagged lines. M.E.P., Z.Z., and W.L. performed data bioinformatics analysis. Z.Z. and W.L. performed ATAC-seq. S.F. performed high-throughput sequencing. S.R. and J.A.W. performed mass spectrometry. M.E.P., I.A., and S.E.J. wrote the manuscript. M.E.P., Z.W., I.A., and S.E.J. coordinated research.

