## Supplementary Materials for "*Arabidopsis* SWR1-associated protein methyl-CpG-binding domain 9 is required for histone H2A.Z deposition"

### Supplementary information:

#### Detailed Bioinformatics Analysis:

#### ChIP-seq analysis.

ChIP-seq fastq reads were aligned to the TAIR10 reference genome excluding chromosome chloroplast and mitochondria with Bowtie[1] using default settings and allowing only uniquely mapping reads. Duplicate reads were removed using SAMtools[2]. Pearson correlation between H2A.Z ChIP-seq replicates was generated with R using normalized (RPM) 1kb binned windows generated using biotoolbox application get_datasets.pl (https://github.com/tjparnell/biotoolbox). Normalized read coverage tracks were generated using the USeq package Sam2Useq application[3]. IGB genome browser was used to visualize the data and to generate snapshots[4]. H2A.Z enriched peaks vs corresponding H3 signal or between mutants for each replicate were determined by callpeak function in MACS2 (v2.1.1.)[5] with the following parameters, -g 1.3e8 –q 0.01 –extsize 200. MBD9, enriched gene were identified similarly as H2A.Z enriched peaks using wild-type Flag ChIP-seq as control. To obtain common H2A.Z enriched peaks between replicates, H2A.Z enriched peaks for each sample were intersected using the USeq package IntersectRegions[3] application with gap =0. Genes corresponding to enriched H2A.Z peaks and MBD9 occupied genes were identified by intersecting with TAIR10 UCSC gene table using the USeq package IntersectRegions application[3]. BioVenn application was used to generate the overlap between H2A.Z enriched genes[6]. ChIP-seq metaplots were generated using NGSPLOT (v4.02.48)[7] with moving window =5. Annotation of peak locations were carried out using the HOMER annotatepeaks function[8]. GO term enrichment was determined using AgriGO[9]. Clustering of H2A.Z profiles in wild-type over MBD9-occupied genes was generated with NGSPLOT (v4.02.48)[7] using *k*-means clustering, n=5, with maximum number of iterations =20 and number of random starts =30.

#### BS-seq analysis.

For generating average methylation plots over MBD9-occupied genes and control set of regions, publicly available wild-type BS-seq data in seedlings was used and analyzed as follows. Trim_galore (http://www.bioinformatics.babraham.ac.uk/projects/trim_galore/) have been used to trim adapters after filtering low quality reads. BS-seq reads were aligned to TAIR10 reference genome by Bismark (v0.18.2)[10] with default settings. Reads with three or more consecutive CHH sites were considered as unconverted reads and have been filtered. DNA methylation levels were defined as #C/ (#C + #T). For meta plot of methylation data, up and down flanking 1000 bp sequences were divided into ten 100bp-bins respectively and MBD9 peaks regions were divided into 10 proportioned bins, then calculate average CG, CHG, and CHH methylation level at these bins.

**RNA-Seq analysis.**
Reads were aligned to TAIR10 using Tophat[11] by allowing up to two mismatches and mapping only to one location. FRPKM values and differential gene expression were analyzed using Cuffdiff[11] with default settings. BioVenn application was used to generate the overlap between differentially expressed genes[6]. Boxplots of expression were generated using R, unpaired two-samples Wilcoxon test was used to determine statistical significance between samples. GO term enrichment was determined using AgriGO[12]. To investigate the misregulation of transcription factors with differentially expressed genes in our mutants, transcriptional factor genes and families were obtained from the Arabidopsis transcription factors database (agris-knowledgebase.org) and intersected with upregulated and down-regulated genes in each mutants. The overlap of transcription factors up-regulated and down-regulated in each mutant was generated using BioVenn.

**Defining H2A.Z gene classes:**

To classify genes according to H2A.Z profiles in wild-type, we used the same approach as described in Coleman-Derr, D. and Zilberman D., 2012. Specifically, H2A.Z levels (RPKM) were obtained for TSS regions (TSS + 500bp) and gene body regions (between TSS +500bp and TES -500bp) for protein coding genes using biotoolbox application get_datasets.pl. We empirically determined that to best represent short genes less than 1500bp, H2A.Z levels were obtained from the entire coding region and the same value was assigned to both TSS and TES region. Next the regions were divided into three groups based on the levels of H2A.Z at gene body (values in log 2 RPKM, low < 3 (9612 genes), medium 3 to <4 (8467 genes), high > =4 (8805 genes)). The three categories were further equally divided into three groups representing low, medium, and high levels of TSS. The categories are designated as L – Low H2A.Z, M- Medium H2A.Z, and H- High H2A.Z for TSS (first letter), and gene body (second letter) as follows (LL, ML, HL, LM, MM, HM, LH, MH, and HH). In total, this analysis represents 26884 protein coding genes. The levels of H2A.Z, gene expression, and gene length are represented by boxplots generated in R for each category of genes. The overlap between each class of genes with H2A.Z and RNA-Seq differential genes for each mutant was determined using BioVenn.

**List of publically available data used in the paper:**
H3K4me3 – 10 day seedlings – (accession number GSE49090)[13]
H3K27me3 – 14 day seedlings – (accession number GSE53620)[14]
DNA methylation - Col-0 seedling BS-seq data (accession number SRR520367)[16]

**List of primers used in the study:**

| Primer Name | Sequence | Description |
|---|---|---|
| JP12457 | CACCGTACATCACATGTTACATGCA | Forward primer ~1kb upstream of MBD9, with CACC tag added for Gateway Cloning into P-ENTR D TOPO (Life Technologies) |

| | | |
|---|---|---|
| JP12458 | GGATCCCTCGGGTTCTTTCCT | Reverse primer -stop codon for MBD9 |
| AtARP6-F | CACCAGCCACAGGAGGAAAGAGAAAT | Forward primer ~1,5kb upstream of ARP6, with CACC tag added for Gateway Cloning into P-ENTR D TOPO (Life Technologies) |
| AtARP6-R | ATGAAAGAATCGTCTACGACACC | Reverse primer -stop codon for ARP6 |
| JP3628 | ttctccaaacgtcgcaacggtctc | FLC RT-PCR |
| JP3629 | gatttgtccagcaggtgacatctc | FLC RT-PCR |
| MAF4-F1 | TCAAGTAACCACCATCACCAACG | MAF4 RT-PCR |
| MAF4-R1 | CAAGAACACACAGAAAAGCACGAA | MAF4 RT-PCR |
| MAF5-F1 | AGACAAAACTCAGGATTATCTTTCACAC | MAF5 RT-PCR |
| MAF5-R1 | ACTTACATTATCCCCTTTTGCTTCTT | MAF5 RT-PCR |
| JP3483 | gatctttgccggaaaacaattggagg | UBQ10 RT-PCR |
| JP3484 | cgacttgtcattagaaagaaagagat | UBQ10 RT-PCR |

### Supplementary Figures

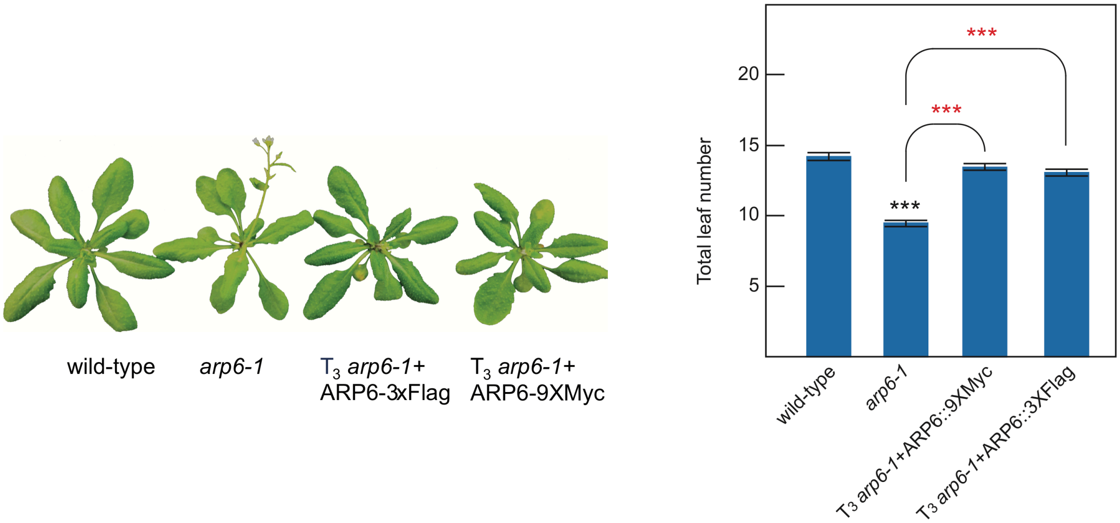


**Supplementary Figure 1. Complementation of the *arp6-1* phenotype.** Morphological phenotype of wild-type, *arp6-1*, $T_3$ *arp6-1+ARP6::3xFlag*, and $T_3$ *arp6-1+ARP6::9xMyc*. Plants were grown for 5 weeks under long-day conditions. Flowering time expressed as the total number of leaves produced by wild-type, *arp6-1*, $T_3$ *arp6-1+ARP6::3xFlag*, and $T_3$ *arp6-1+ARP6::9xMyc* from 16 plants ± standard deviations under the same conditions is also shown. Paired two-tailed Student's t-test was used to determine significance between wild-type and the mutant (black asterisks) or the transgenic lines (red asterisks); ns p-value > 0.05, * p-value <= 0.05, ** p-value <= 0.01, *** p-value <= 0.001.

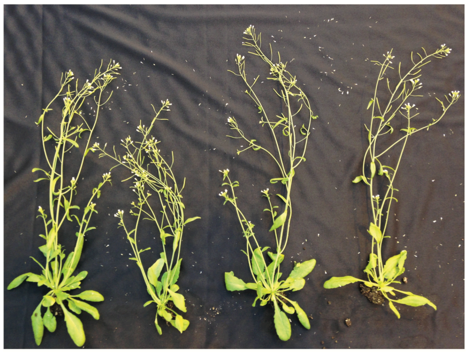


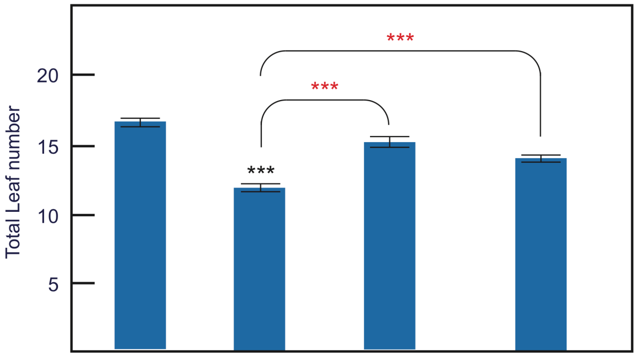


**Supplementary Figure 2. Complementation of the *mbd9-3* phenotype.** Morphological phenotype of wild-type, *mbd9-3*, $T_4$ *mbd9-3+MBD9::3xFlag*, and $T_4$ *mbd9-3+MBD9::9xMyc*. Plants were grown for 7 weeks under long-day conditions. Flowering time expressed as the total number of leaves produced by wild-type, *mbd9-3*, $T_4$ *mbd9-3+MBD9::3xFlag*, and $T_4$ *mbd9-3+MBD9::9xMyc* from 12 plants ± standard deviations at 5 weeks is shown. Paired two-tailed Student's t-test was used to determine significance between wild-type and the mutant (black asterisks) or the transgenic lines (red asterisks); ns p-value > 0.05, * p-value <= 0.05, ** p-value <= 0.01, *** p-value <= 0.001.

a

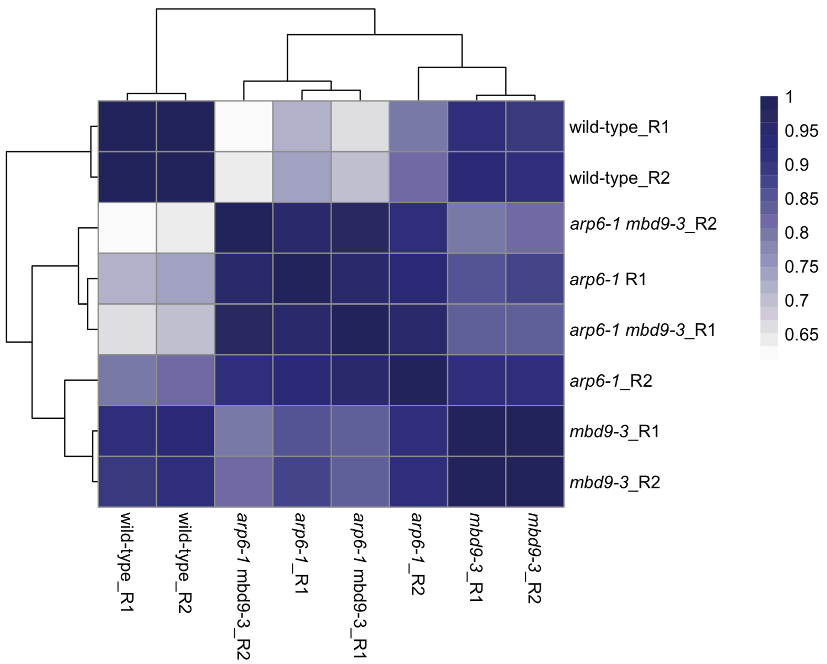

wild-type_R1
wild-type_R2
arp6-1 mbd9-3_R2
arp6-1 R1
arp6-1 mbd9-3_R1
arp6-1_R2
mbd9-3_R1
mbd9-3_R2
1
0.95
0.9
0.85
0.8
0.75
0.7
0.65
wild-type_R1
wild-type_R2
arp6-1 mbd9-3_R2
arp6-1_R1
arp6-1 mbd9-3_R1
arp6-1_R2
mbd9-3_R1
mbd9-3_R2


b

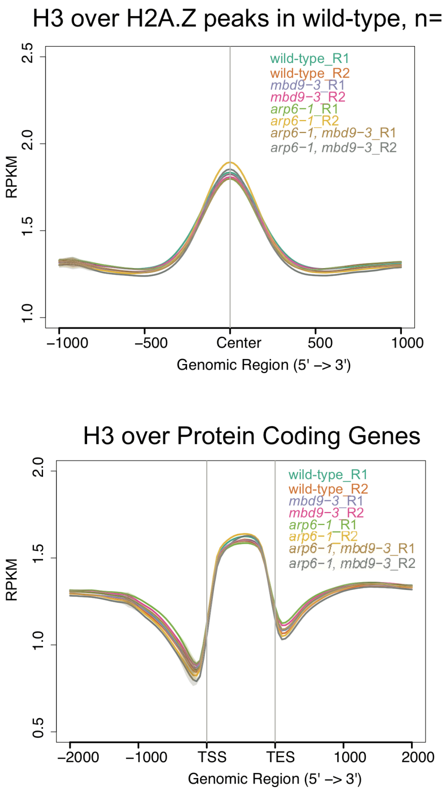

H3 over H2A.Z peaks in wild-type, n=18481
wild-type_R1
wild-type_R2
mbd9−3_R1
mbd9−3_R2
arp6−1_R1
arp6−1_R2
arp6−1, mbd9−3_R1
arp6−1, mbd9−3_R2
RPKM
1.0
1.5
2.0
2.5
−1000
−500
Center
500
1000
Genomic Region (5' −> 3')
H3 over Protein Coding Genes
wild-type_R1
wild-type_R2
mbd9−3_R1
mbd9−3_R2
arp6−1_R1
arp6−1_R2
arp6−1, mbd9−3_R1
arp6−1, mbd9−3_R2
RPKM
0.5
1.0
1.5
2.0
−2000
−1000
TSS
TES
1000
2000
Genomic Region (5' −> 3')


c

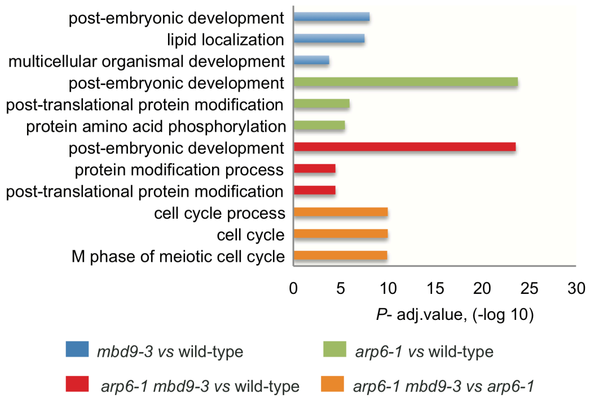

post-embryonic development
lipid localization
multicellular organismal development
post-embryonic development
post-translational protein modification
protein amino acid phosphorylation
post-embryonic development
protein modification process
post-translational protein modification
cell cycle process
cell cycle
M phase of meiotic cell cycle
0
5
10
15
20
25
30
P- adj.value, (-log 10)
mbd9-3 vs wild-type
arp6-1 vs wild-type
arp6-1 mbd9-3 vs wild-type
arp6-1 mbd9-3 vs arp6-1

**Supplementary Figure 3. Correlation of H2A.Z ChIP-seq and profiles of H3 ChIP-seq replicate data. (a)** Pearson correlation of normalized 1 kb binned H2A.Z ChIP-seq signal (RPM) for indicated replicates. **(b)** Distribution of normalized H3 ChIP-seq signal (RPKM) for each ChIP-seq replicate over H2A.Z common peaks in wild-type and protein-coding genes. **(c)** GO term analysis for H2A.Z-depleted genes (macs2 peak caller, q-value less than 0.01). *P*-adjusted value in –log(10) is shown for top three GO classes for the indicated mutants.

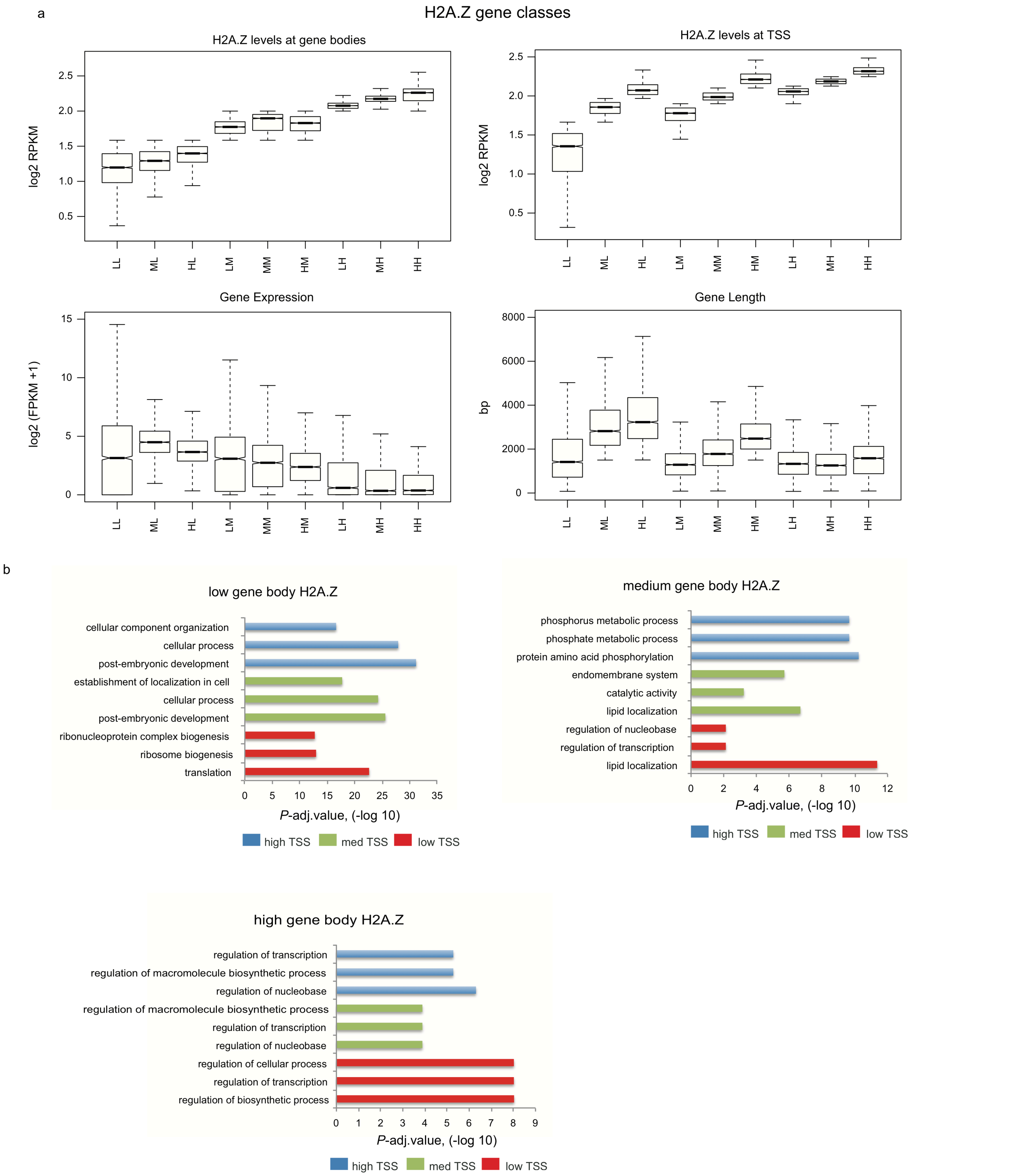


**Supplementary Figure 4. Features of gene classes characterized according to their levels of H2A.Z at TSS and gene body.** H2A.Z levels (RPKM) were calculated for TSS regions (TSS + 500 bp) and gene body regions (between TSS +500 bp and

TES -500 bp) for protein-coding genes. **(a)** Notched boxplots of H2A.Z levels (RPKM) at gene bodies and TSS from merged replicates of H2A.Z ChIP-seq in wild-type. Outliers not plotted. Gene expression in $\log_2$ (FPKM+1) in wild-type and gene length in bp for each class. Labels H – high, M – medium, L- low, refer to levels of H2A.Z at TSS (first letter) and gene body (second letter). (b) GO term analysis for the nine classes of genes. *P*-adjusted value in –log(10) is shown for the top three GO classes for the indicated mutants.

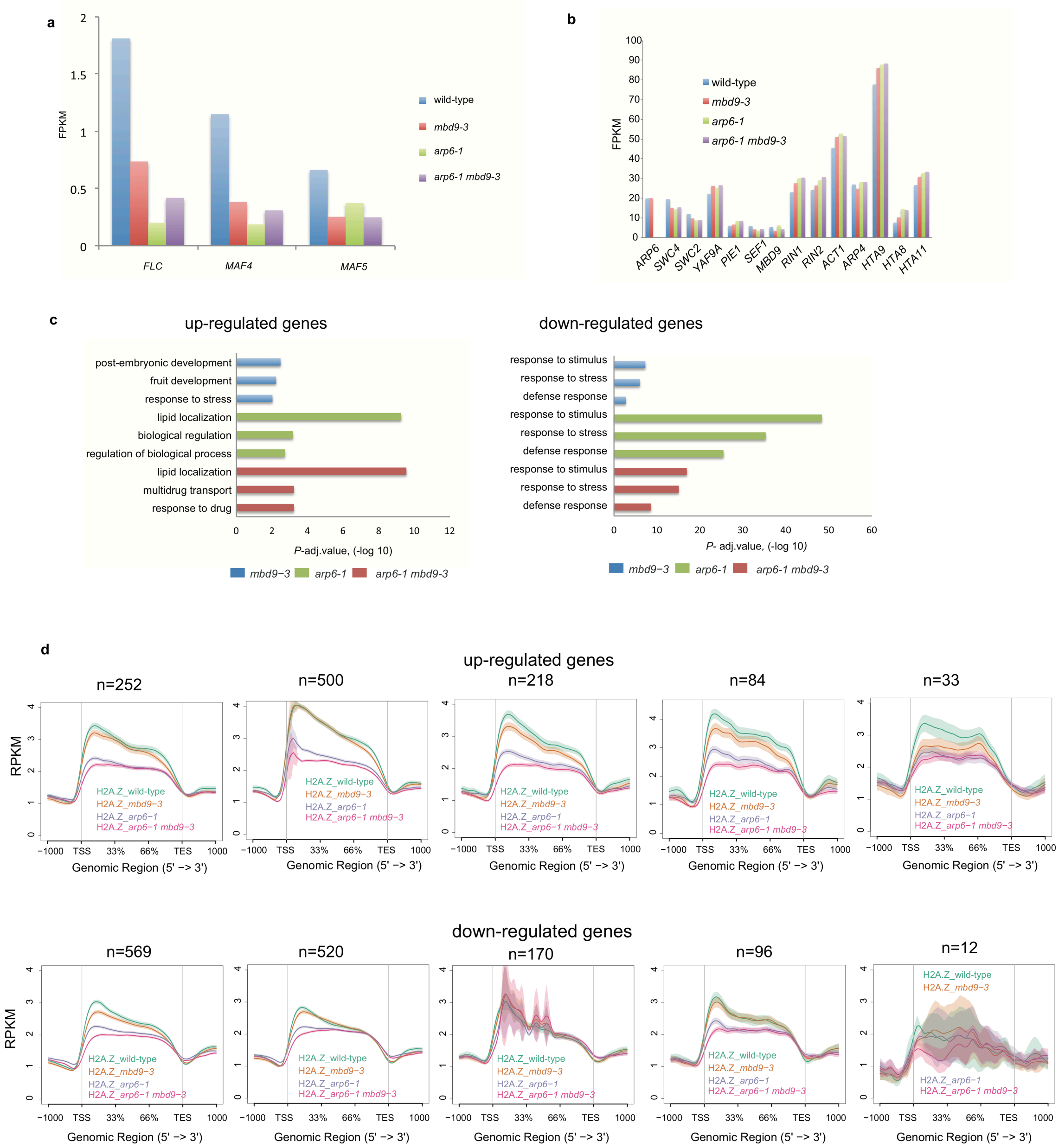


**Supplementary Figure 5. (a)** Normalized expression levels (FPKM) of *FLC, MAF5,* and *MAF4* from RNA-Seq (FPKM) in wild-type *mbd9-3, arp6-1*, and *arp6-1 mbd9-3* from four independent replicates for each sample. **(b)** Normalized expression levels (FPKM) of SWR1 complex components and H2A.Z genes in wild-type, *mbd9-3, arp6-1*, and *arp6-1 mbd9-3* from four independent replicates for each sample. **(c)** GO term analysis for significantly up-regulated and down-regulated genes in each mutant vs wild-type control. *P*-adjusted value in –log(10) is shown for the top three GO classes for each mutant. **(d)** Distribution of normalized H2A.Z ChIP-seq signal (RPKM) from merged replicates over classes of overlapping up-regulated and down-regulated genes based on RNA-Seq in wild-type, *mbd9-3, arp6-1*, and *arp6-1 mbd9-3*.

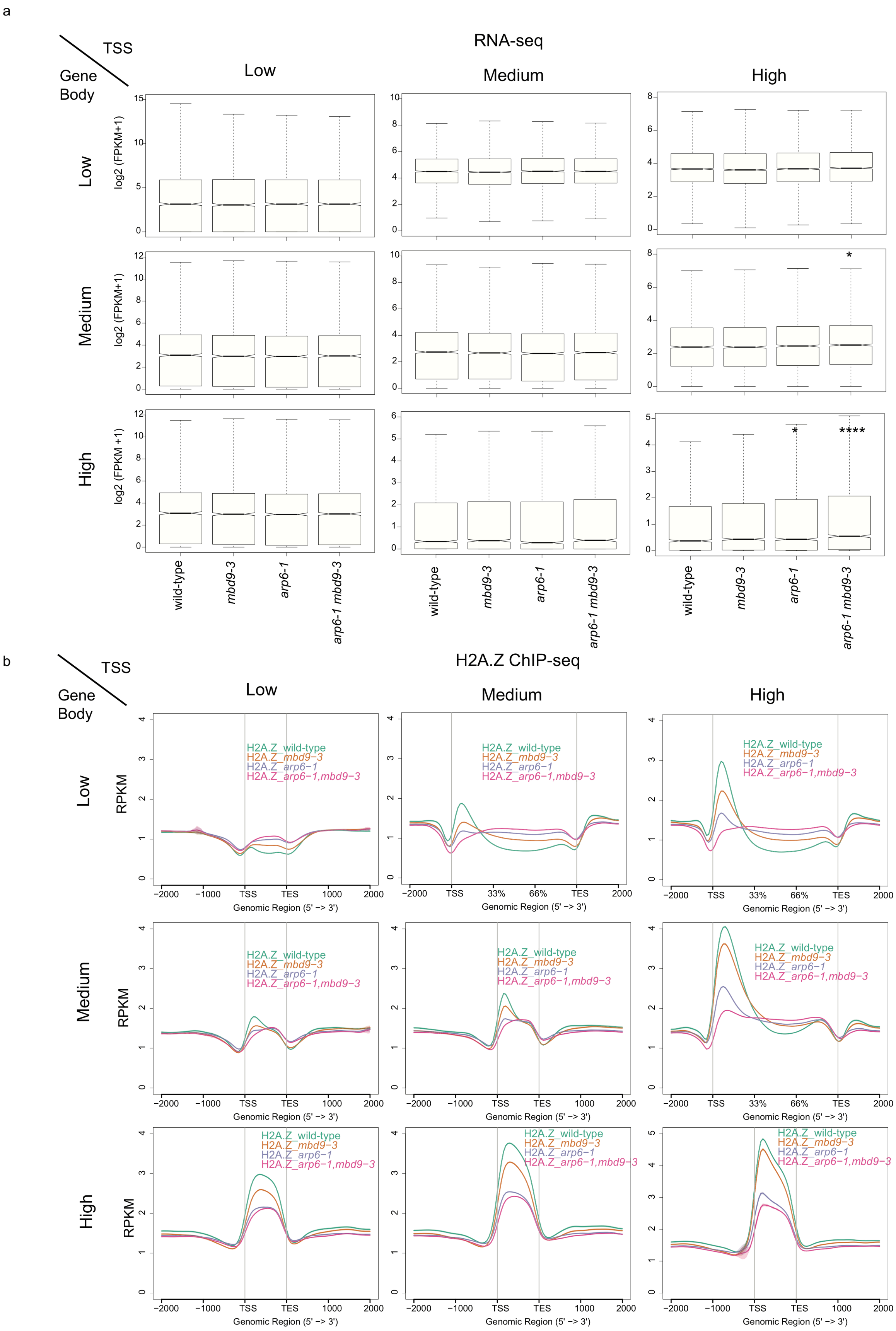
a
TSS
Gene Body
RNA-seq
Low
Medium
High
log2 (FPKM+1)
log2 (FPKM +1)
*
****
wild-type
mbd9-3
arp6-1
arp6-1 mbd9-3
b
TSS
Gene Body
H2A.Z ChIP-seq
Low
Medium
High
RPKM
H2A.Z_wild-type
H2A.Z_mbd9−3
H2A.Z_arp6−1
H2A.Z_arp6−1,mbd9−3
−2000
−1000
TSS
TES
1000
2000
33%
66%
Genomic Region (5' −> 3')

**Supplementary Figure 6.** Expression and H2A.Z profiles at nine classes of H2A.Z-occupied genes in wild-type, *mbd9-3, arp6-1*, and *arp6-1 mbd9-3.* **(a)** Notched boxplots of average normalized RNA-Seq reads $\log_2$ (FPKM +1) for nine classes of H2A.Z genes according to Supplementary Fig. 4. Outliers not plotted. Unpaired two-samples Wilcoxon test was used to determine significance between wild-type and mutants, only significant values are shown; * p-value <= 0.05, ** p-value <= 0.01, *** p-value <= 0.001, **** p-value <= 0.0001. **(b)** Distribution of normalized H2A.Z ChIP-seq signal (RPKM) from merged replicates over the nine classes of H2A.Z genes according to Supplementary Fig. 4.

Transcription factors up and down regulated in SWR1 mutants

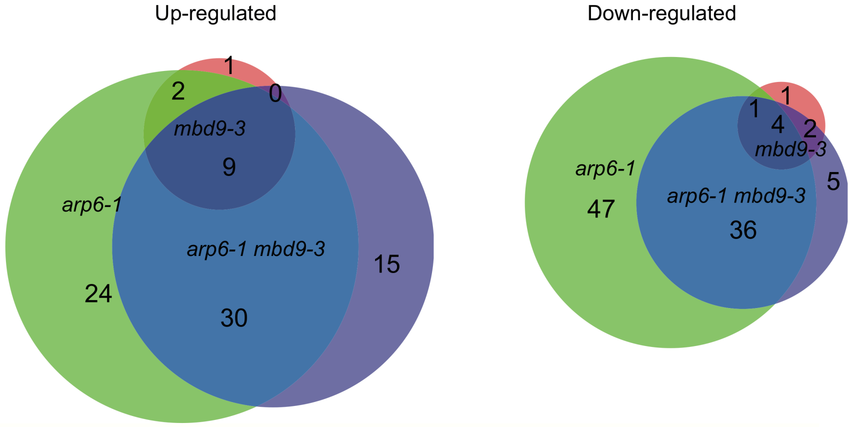


Transcription factor families up-regulated in SWR1 mutants

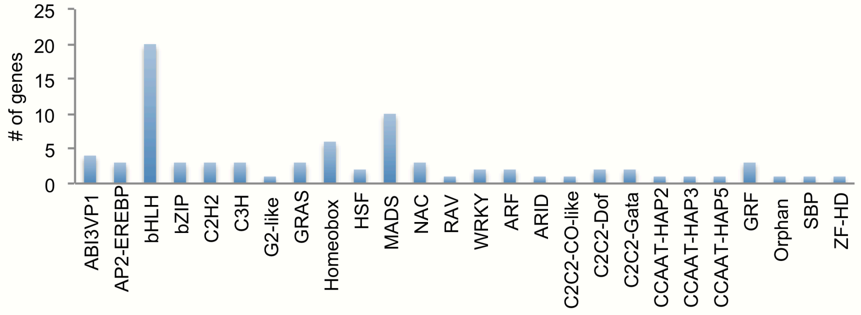


Transcription factor families down-regulated in SWR1 mutants

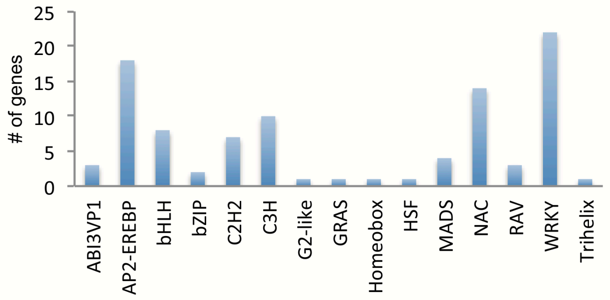

**Supplementary Figure 7.** Intersection of significantly up-regulated and down-regulated genes with transcription factors obtained from AtTFDB (agris-knowledge.com). Distribution of transcription factor families that intersected with up-regulated and down-regulated genes in wild-type, *mbd9-3, arp6-1*, and *arp6-1 mbd9-3.*

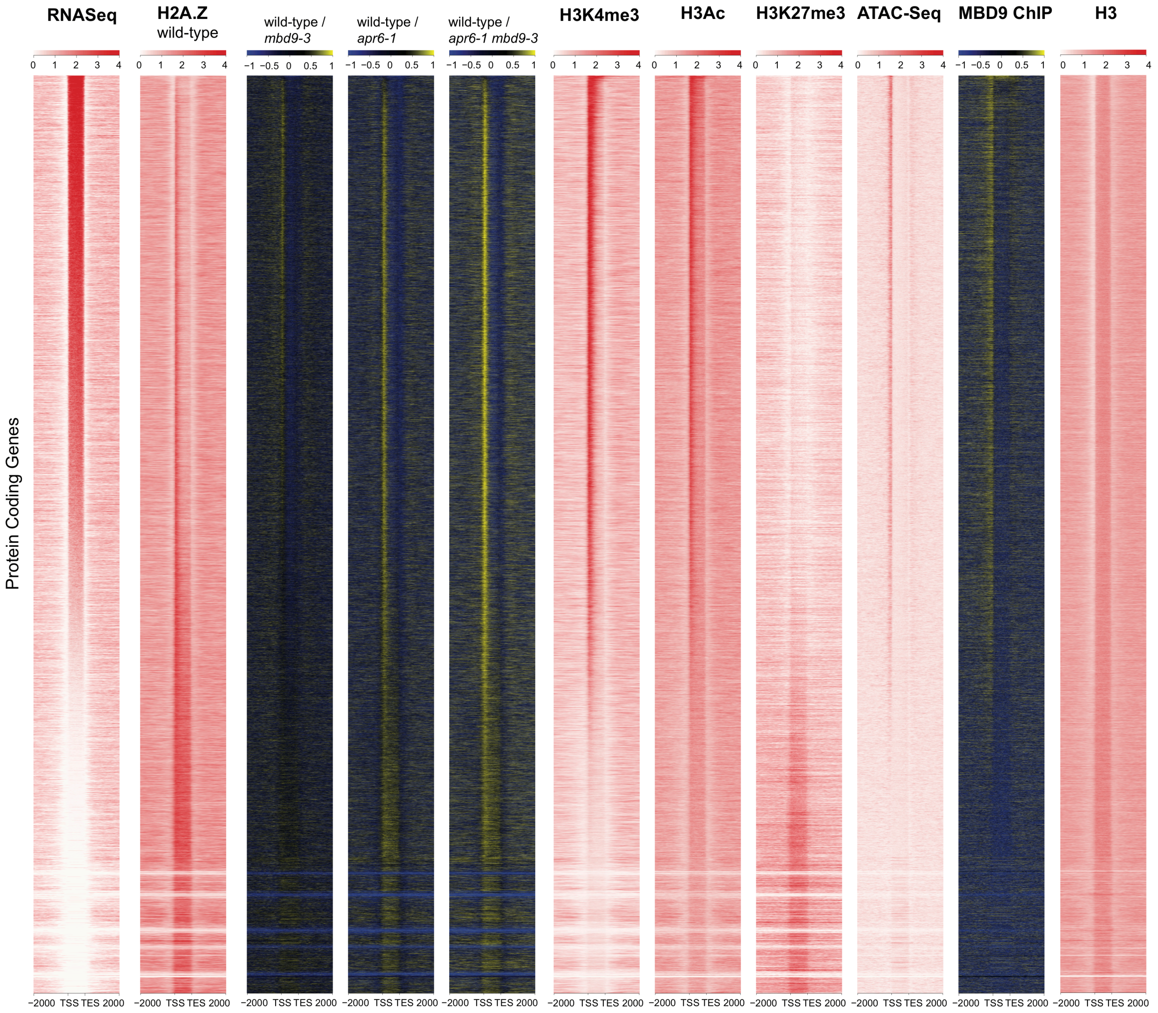


**Supplementary Figure 8.** Heatmap of normalized expression (RPKM) and histone modifications (RPKM) plotted in relation to decreasing levels of expression in wild-type for RNA-Seq, H2A.Z in wild-type, ratio plots for H2A.Z vs wild-type for *mbd9-3, arp6-1,*

*arp6-1 mbd9-3* mutants, H3K4me3, H3AC, H3K27me3, ATAC-Seq, MBD9_ChIPSeq, and H3 in wild-type.
